# Global distribution patterns of an ocean-surface dwelling animal are associated with organismal mirror asymmetry

**DOI:** 10.1101/2025.11.04.686608

**Authors:** Tom Iwanicki, Nikolai Maximenko, Casey Dunn, Samuel Church, Deb Goodwin, Rebecca R. Helm

## Abstract

Organismal mirror asymmetry is common in nature, where an individual may have either a left- or right-handed form; however, the causes and consequences of this asymmetry have long fascinated and perplexed biologists. The cnidarian by-the-wind sailor jelly *Velella* is globally distributed, floating on the sea surface and harnessing the wind with a protruding fleshy sails that develop either a left-handed or right-handed orientation. For over 60 years, scientists have postulated that this handedness is related to wind-sorting, and enables a broad and consistent distribution of handedness types. In this hypothesis, due to prevailing winds, left handed sailors are more likely to wash ashore in the northern hemisphere, while right handed sailors are more likely to occur in central gyres. The opposite pattern is predicted in the southern hemisphere. Here, we leverage the global reach of community (citizen) science, genetics, and oceanographic modeling, to examine this enduring hypothesis. Our results are consistent with predicted patterns. Proportionally more left-handed sailors washed ashore in the Northern Hemisphere and right-handed sailors in the Southern Hemisphere. When sampling through the central part of the North Pacific Subtropical Gyre, we found almost exclusively right-handed sailors, whereas left-handed sailors occurred on the edge of the gyre toward the coast. These observations are concordant with calculated mean streamlines that would distribute left- and right-handed sailors in different patterns. Some authors suggest these handedness types are different species, consistent with their broad differences in distribution. However, we found that both left- and right-handed sailors in the western North Atlantic form a single population, with no detectable genomic differentiation or evolutionary structure in a mitochondrial phylogeny. We next examined the drivers of this differential distribution. At a regional scale, the mixed population off the western coast of Portugal experienced episodic strandings of left- and right-handed sailors in proportions correlating with observed wind and modeled sailing vectors, further supporting wind sorting of sail types as the primary cause of differential distributions. Modeling suggests that global dispersal is dependent on mirror asymmetry. Left-handed sailors are more likely to cross the equator from north-to-south, while right handed sailors are more likely to cross from south-to-north. Combined, our data provides evidence that mirror asymmetry is an important factor in a species global distribution.

## Summary and Background

For over 100 years, scientists have puzzled over why some species exhibit dimorphic asymmetry (Levin and Palmer 2007; Palmer 1996; Palmer 2005; Palmer 2004). This asymmetry presents as “mirroring” or differences in handedness, with left- or right-handed animals co-occuring between closely related species, or even within populations. Yet the causes and consequences of asymmetry are not well understood. Here, we investigate a decades-old hypothesis on asymmetry in a surface-dwelling hydrozoan. By-the-wind sailor (*Velella;* Fig. 1A) harnesses the wind with a fleshy rigid sail. The sail grows at an angle, either to the left or right, of the main axis. This asymmetry begins in early development and is fixed throughout the animal’s life (Woltereck, 1904). By-the-wind sailor are globally distributed and can aggregate on the sea surface by the tens- or hundreds of thousands (Fig. 1B). Bieri reviewed global observations and noted stark differences in left- and right-handed distribution, but could not determine the cause (Bieri, 1959). Some speculated that left- and right-handed forms were species characteristics (Eschscholtz 1829). In contrast, in 1958, Soviet Scientist A.I. Savilov hypothesized that *Velella* is a single species, and that differential dispersal was associated with this dimorphic sail angle (Savilov 1958,1969). *Velella* sail downwind at an angle approximately 30-60 degrees relative to the prevailing wind direction, with left-handed sailors traveling to the left and right-handed sailors traveling to the right (Fig. 1C) (Mackie 1962). Savilov’s hypothesis predicts that in the Northern Hemisphere, left-handed sailors are blown toward the coastal margins and right-handed sailors are concentrated inside oceanic gyres. In the Southern Hemisphere, the opposite pattern holds true, with right-handed sailors occurring in greater abundance along the coastline and left-handed sailors concentrating inside oceanic gyres (Fig. 1D).

**Fig. 1.**
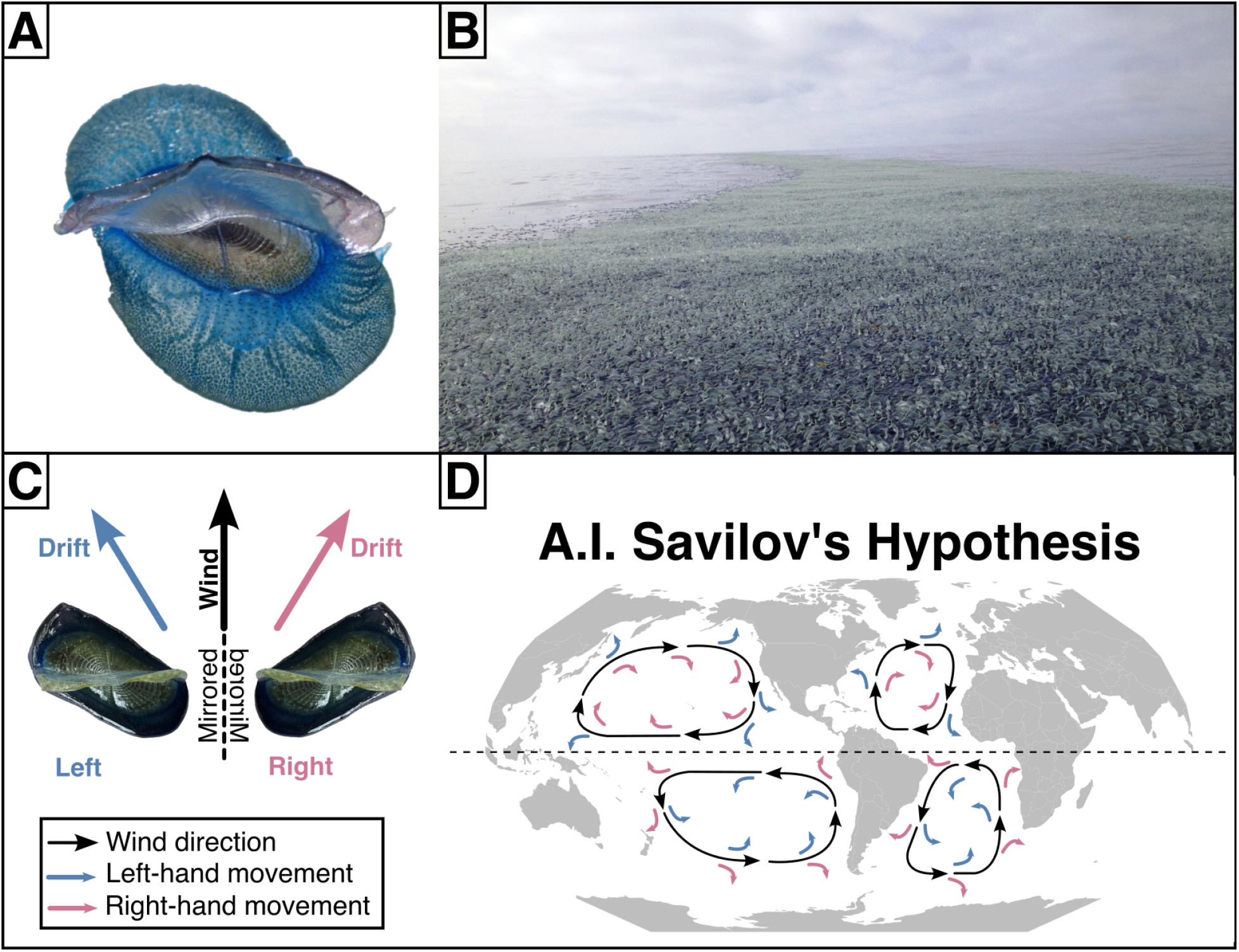
A: Photograph of a left-handed By-the-wind sailor (*Velella*). B: Large raft of thousands of by-the-wind sailor floating on the sea surface viewed from a boat off the coast of California (photo credit: Scott Horton). C: Posture of By-the-wind sailor of different handedness and their drift velocity vectors relative to the wind direction, derived from Mackie (1962). D: A simplified map summarizing and extrapolating globally A.I. Savilov’s hypothesis for why and where we observe dimorphic sail direction, arrows show the prevailing wind direction and fate of left- and right-handed sailors.

Understanding how asymmetry relates to distribution is critical for understanding the broader ocean’s surface. *Velella* is an integral component of the surface neustonic ecosystem–the largest contiguous habitat on Earth, and one of two key genera at the ocean’s surface to exhibit directional asymmetry (including *Physalia* spp.) *Velella* can form massive aggregations at the air-sea interface, completely covering the surface for dozens of kilometers (Helm 2021). Likewise, this species (along with *Physalia*) is a critical food source and habitat for dozens of species (Savilov 1956, Helm 2021a,b).

If Savilov’s hypothesis is correct, dimorphism may be an important component of the broad dispersal of this species. However, no global study exists examining *Velella* distribution. In one of the most up-to-date accounts, R. Bieri concluded that for lack of data across seasons and size classes “…*it is apparent that the origin and occurrence of the different sizes and morphological forms of Velella and Physalia are not yet satisfactorily explained*.” (Bieri, 1959).

Here, we leverage participatory science observations, phylogenetics, population genomics, and surface ocean transport modeling to examine the role of handedness in distribution.

## Results & Discussion

### Community science

We combined 11,115 iNaturalist observations processed by 1,169 volunteers on the crowd-source platform Zooniverse (Iwanicki & Helm 2025) and 67 plankton tows from two sea-going expeditions with professional and community scientists to quantify the occurrence of left- and right-handed sailors from around the world. The iNaturalist observations covered the global extent of *Velella* distribution, with several regions exhibiting greatest sampling effort, namely, the west coast of North America, the Mediterranean, and the coasts of Southern Australia and New Zealand. Zooniverse volunteers assessed iNaturalist observations and assigned the count, condition (i.e., dead or alive), and sail direction (i.e., left or right) resulting in high quality data following validation as described in Iwanicki & Helm (2025) (Supp. File A).

In total, 6,421 observations were confidently assigned sail direction with proportionally more left-handed sailors (n = 5,000) than right-handed sailors (n = 438) in the Northern Hemisphere, and slightly more right-handed sailors (n = 540) than left-handed (n = 443) in the Southern Hemisphere (X-squared = 1417, p-value < 0.0005) (Fig. 2A). Additionally, left- and right-handed sailors appear within each hemisphere in proportions significantly different from a 50:50 distribution (Northern Hemisphere: X-squared = 3827, p < 0.0005; Southern Hemisphere: X-squared = 9.57, p-value < 0.01).

**Fig. 2.**
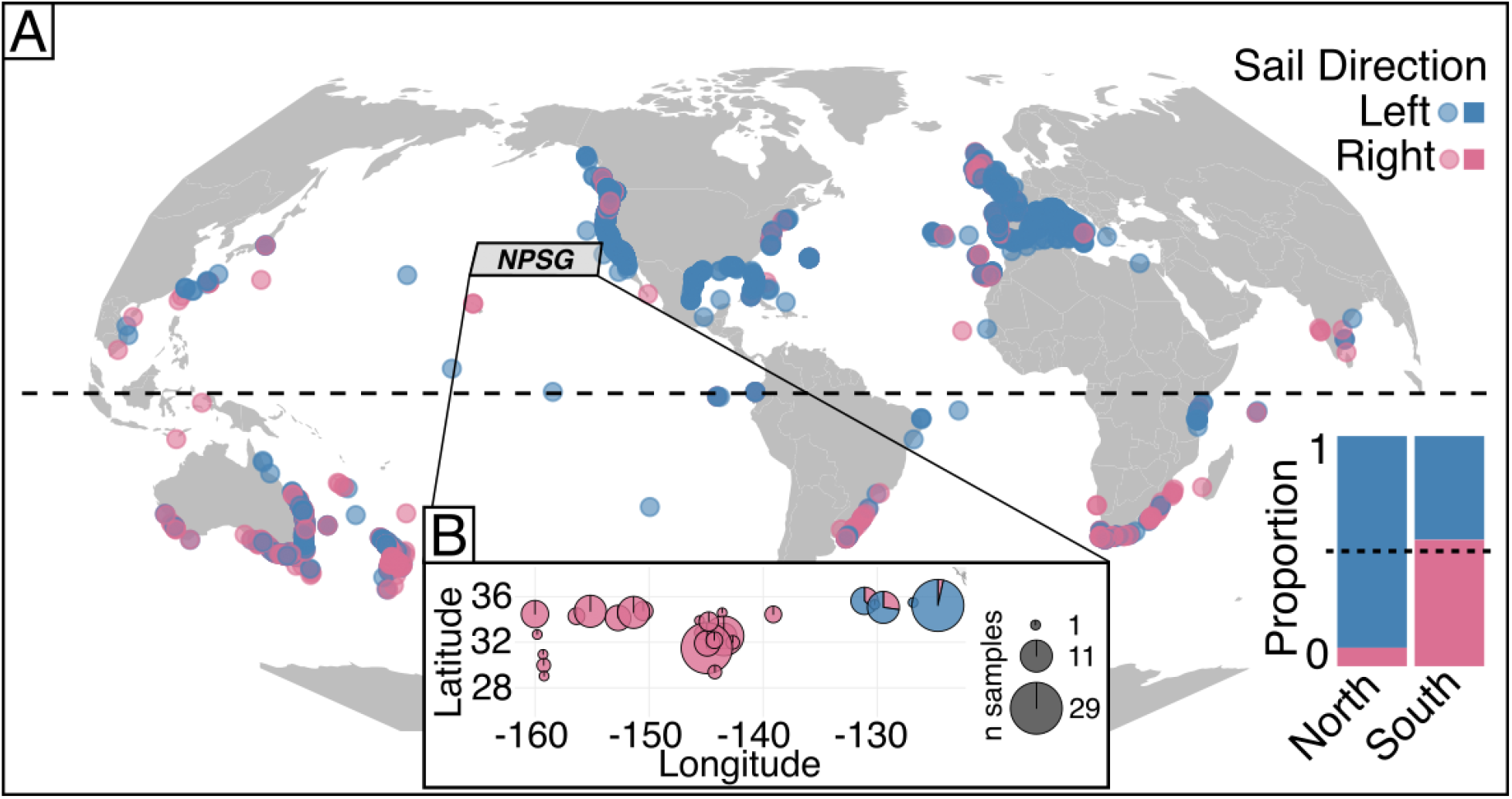
A: Map of iNaturalist observations of *Velella velella* with the sail direction determined by volunteers on the community science platform Zooniverse (Iwanicki & Helm 2025). B: pie-charts showing the proportion of left (blue) and right (red) *Velella velella* observations plotted to the location from two sea-going community science expeditions (SEA and The Vortex Swim) to the eastern North Pacific Subtropical Gyre (marked as NPSG).

We collected at-sea observations through the North Pacific subtropical gyre, where we would expect more right-handed sailors than left-handed sailors, with its center oscillating around 32°N,145°W (Moore et al. 2001). The first expedition occurred between June and August 2019 and involved neuston net tows performed by the sailing crew accompanying long-distance swimmer Benoît Lecomte (https://benlecomte.com/; The Vortex Swim). Chong *et al*. (2023) previously reported patterns of total plastic and neuston abundance in the gyre using these data. The second expedition was aboard the Sea Education Association (SEA) cruise between June and July 2022 and used the same collection protocol as The Vortex Swim. From photographs of filtered neuston net tows, we collected a total of 314 *Velella* observations, of which we were able to assign sail direction to 43 left-hand, 133 right-hand across 27 stations (Supp. File B).

Right-handed sailors were dominant in the central part of the gyre and left-handed sailors appeared toward the west coast of North America and in proximity of the California Current (Fig. 2B).

### Molecular analyses

Given the broad differences in left- and right-handed distributions, we examined if there were genetic differences in left- and right-handed *Velella* where they co-occur. Eschscholtz (1829) hypothesized that left- and right-handed dimorphic forms are species characteristics. This was later rejected by Chun (1897) based on their other similarity in morphology. Later, the observation that predominantly left-handed sailors occur in the Mediterranean, which was also observed in our community science data, led Moser (1925) to predict genetic isolation with each form. We performed genome skimming and assembled mitochondrial genomes for phylogenetics and mapped reads to a *Velella* transcriptome for population genetic analyses for samples collected from the United States Pacific and Atlantic coasts.

Genomic DNA was sequenced from 24 left-hand and 24-right-hand *Velella* from the Atlantic that co-occurred on the coast of North Carolina in October 2024 (Fig. 3A), and 12 left-hand *Velella* from the Pacific near Monterey Bay, California. Molecular analyses were performed following methodology detailed in Church et al. 2025. We estimated the *Velella* nuclear genome size to be 3.16 Gb, with 2.63% heterozygosity at 21.2x coverage with a best fit model at 97.1%. Given the large nuclear genome size and high degree of heterozygosity, we proceeded with population genetic analyses using a publicly available transcriptome rather than assembling a *de novo* genome. This transcriptome mapped approach can be robust for inferring population statistics compared to genome mapped reads as demonstrated by Church et al. (2025). Gene trees were used to deduplicate transcripts (Guang et al. 2021, Munro et al. 2022), leaving 23,793 transcripts. Reads mapped to 19,511 transcripts containing 29,391 windows and 892,458 SNPs which revealed no population structure among left- and right-handed sailors in the Atlantic with a pairwise *Fst* of effectively zero (Kruskal-Wallis chi-squared = 5759.53, df = 2, p-value < 0.005) (Fig. 3B).

**Fig. 3.**
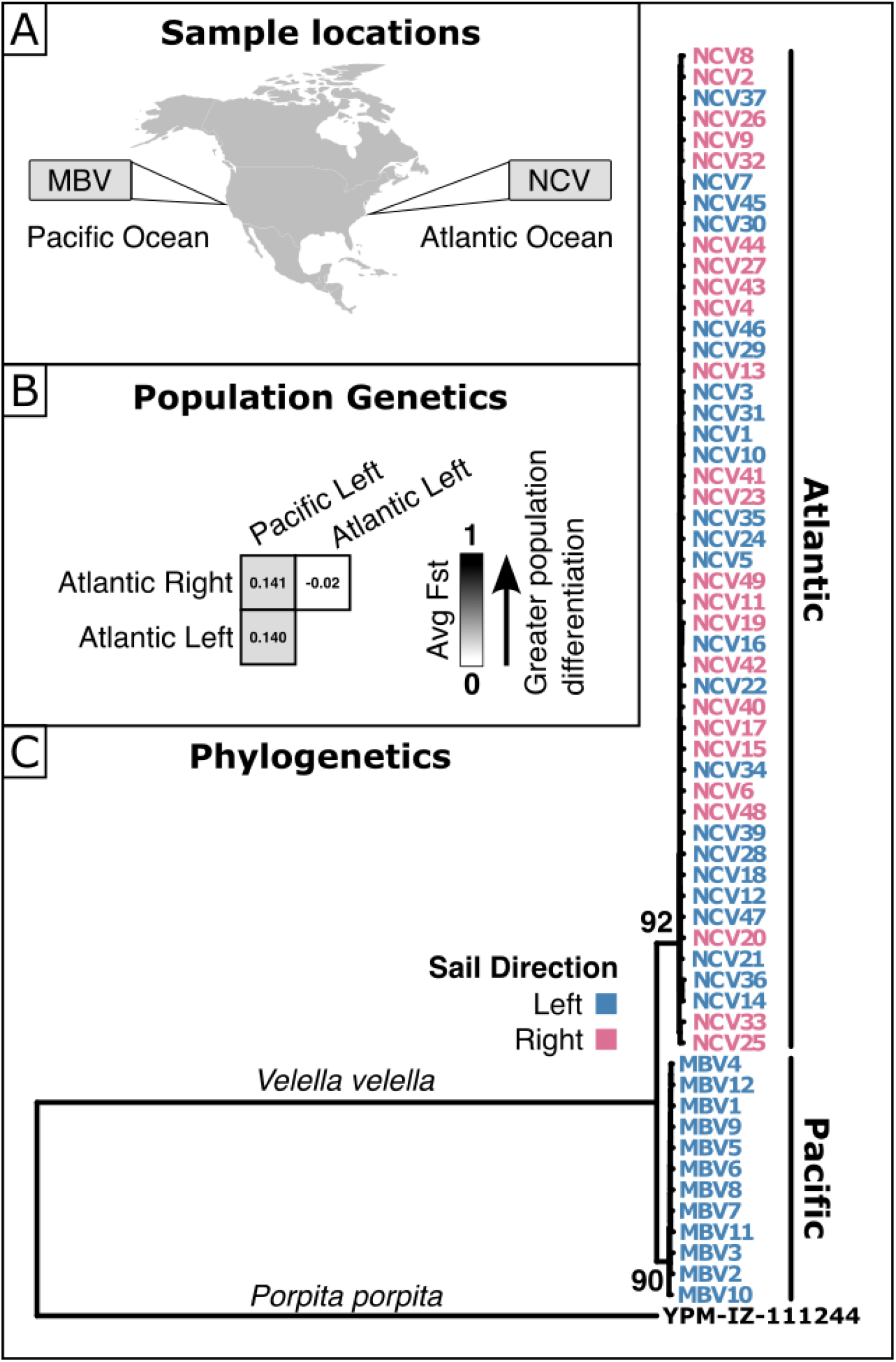
Molecular analyses of *Velella*. A: Map of *Velella* sample collections from Monterey Bay, California (MBV) and North Carolina (NCV). B: Genomic reads mapped to a transcriptome derived pair-wise population differentiation estimate (*Fst*) reveals no population differentiation among left- and right-handed *Velella*, and moderate differentiation among Pacific and Atlantic populations. C: Maximum likelihood consensus tree derived from IQTREE using an alignment of whole mitochondrial genomes from *Velella* and outgroup *Porpita* (Yale Peabody Museum Catalog number: YPM-IZ-111244). Tip label color corresponds to sail direction: blue = left-handed, red = right-handed, black = outgroup.

Atlantic and Pacific *Velella* had moderate population differentiation, which was not different among left- and right-handed sailors (*Fst* = 0.140 and 0.141, respectively; Dunn’s test p = 0.2867). Mitochondrial genomes were assembled, aligned, and a phylogenetic tree was inferred that revealed no pattern among the sail direction (Fig. 3C) (Supp. File C).

Using both population genetic and phylogenetic analyses, we found no evidence that left- and right-handed *Velella* are part of distinct isolated populations or separate species within the western North Atlantic. Instead, left and right handedness occurred within the same populations.

### Oceanographic analyses

Savilov hypothesized that differences in distribution of dimorphic forms is related to wind sorting. To test this, we examined wind drift using an ocean drift model (Maximenko and Hafner 2024), which was developed and used to simulate the drift of various types of marine debris and validated using observational reports (Maximenko et al. 2018; Berg et al. 2024).

We first used observations of Mackie (1962) on left-handed *Velella* drift angle and speed to determine how it responds to known wind and extrapolated right-handed sailors by flipping the angle sign (Fig. 1C). We then applied these wind coefficients to long-term averaged wind vectors and added model mean surface currents to derive streamlines for left- and right-handed *Velella* (Fig. 4A,B). While the mean streamlines includes more complexity than Savilov’s simplified model, it clearly confirms pathways of right- and left-handed *Velella* converging towards the subtropics in the Northern and Southern Hemisphere, respectively.

**Fig. 4.**
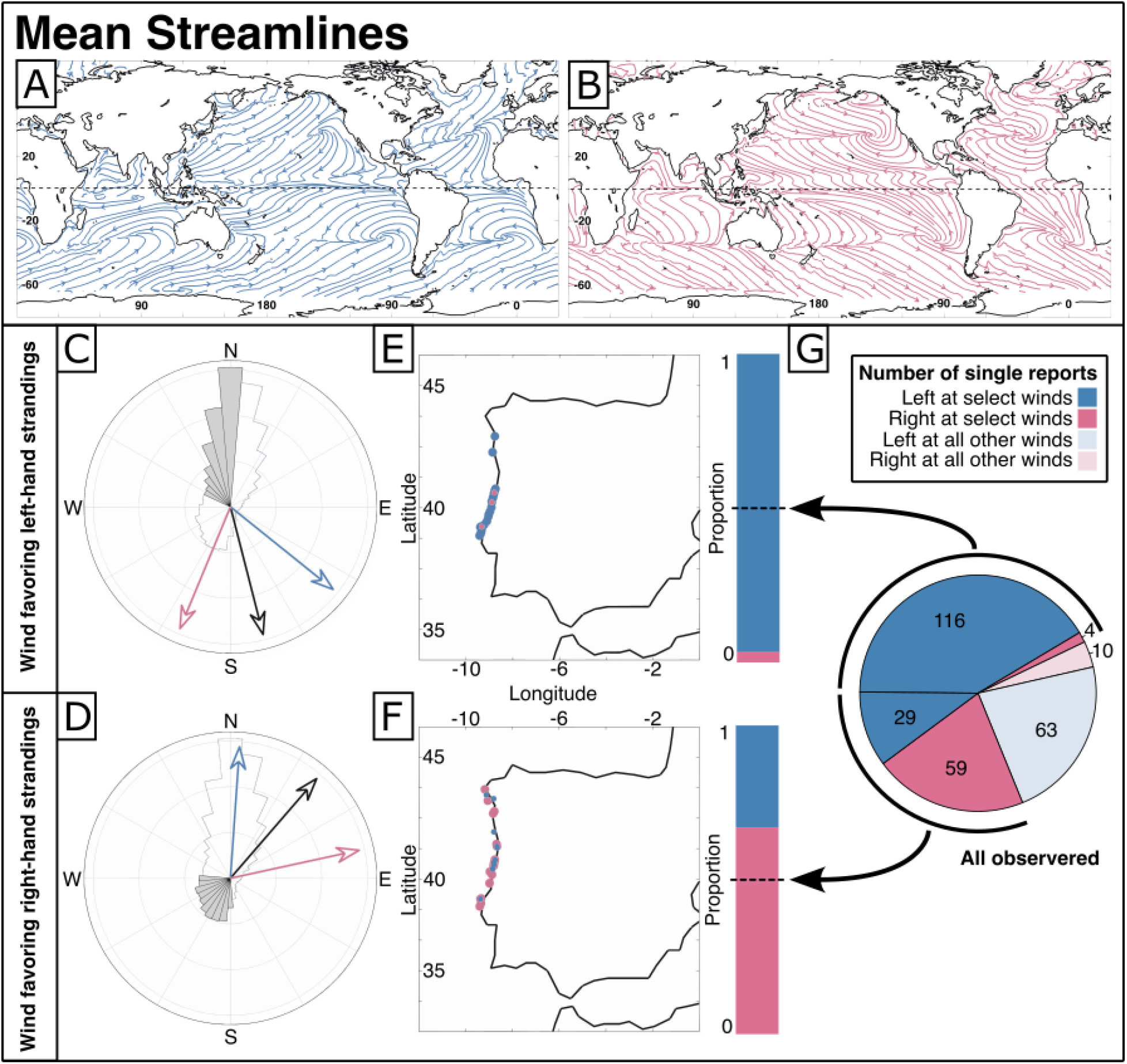
Dependence of *Vellela* composition in reports from the Portugal west coast from wind direction. A,B: Mean simulated streamlines for *Velella* (left-handed: blue; right-handed: red). C,D: mean wind roses, wind directions selected for the subset (black sectors and vector) and associated velocity vectors of left-handed (blue) and right-handed (red) *Velella*. E,F: The coastline and locations of left-handed (blue) and right-handed (red) reports. Locations of reports of dominant handedness are shown with larger dots placed in the bottom layer. G: Pie diagram and stack barplots with proportions summarizes composition of *Velella* reports during winds favoring beaching of left-handed (E) and right-handed (F) sailors.

Additionally, the model suggests that global dispersal is dependent on mirror asymmetry. The north-to-south dispersal of *Velella* across the equator is dominated by left-handed sailors (Fig 4A). In contrast, the south-to-north transport of *Velella* across the equator dominated by right-handed sailors (Fig 4B).

We next examined how well our model performs relative to observed data. We used iNaturalist reports from a region (western Portugal) with highly variable winds that is dominated by one morph (left-hand) but may experience episodic strandings of the other morph (right-hand). To determine the role that wind may play in these episodic strandings, we examined 281 reports. A majority of these reports (n=208) were left-handed, while a subset (n = 73) were right-handed. Left- and right-handed *Velella* arrive under different wind conditions (Fig. 4C-G). Under northerly-westernortherly winds (Fig. 4C), left-handed *Velella* (blue arrow) drift toward shore, while right-handed *Velella* (red arrow) exhibit a long-shore velocity, resulting in observations of 116 left-handed *Velella* stranding, and only 4 right handed *Velella*. In contrast, southwesterly winds (Fig. 4D) induce onshore flux of right-handed *Velella*, and longshore transport of left-handed *Velella*, with only 29 left-handed *Velella* reported compared to 59 right-handed *Velella*. The pie chart and stacked barplots (Fig. 4G) show the proportion of *Velella* that are actually stranded under select wind conditions as dark-shaded wedges, while strandings at all other winds are light-shaded. These results support our broader model findings that wind is a critical real-world driver of left- and right-handed distribution.

### Modern approaches for addressing an old question

The cause and consequence of organismal asymmetry have been a subject of study for hundreds of years. Here, we find support for A.I. Savilov’s decades-old hypothesis that *Velella* global distribution is correlated with its sail mirror asymmetry. We report that i) left-and right-handed sailors washed ashore in greater proportions in the Northern and Southern Hemispheres, respectively, ii) right-handed sailors were observed in greater abundance inside the North Pacific Subtropical Gyre, iii) no population differentiation between left- and right-handed sailors was detected in the western North Atlantic, and iv) that winds facilitate transport of asymmetric forms to disparate ocean regions, including differential crossing of the equator.

Combined, these data demonstrate that an asymmetric trait may be a key driver of the observed global distribution in this species.

Still, many questions remain unanswered. We do not know the causal mechanism of this asymmetry. Is it genetically determined (directional asymmetry), or randomly determined (antisymmetry; Palmer, 2004)? The genetics of asymmetry have important implications for how this feature is maintained. In handedness under balancing selection? Or is it random? Future studies are needed to determine the genetic and developmental basis of asymmetry. Asymmetry appears to facilitate global population connectivity across ocean basins. Future studies should examine the global population structure and connectivity of this species in relation to sail form, and the sailless sister-species (of a monotypic genus) *Porpita porpita*. Finally, *Velella* is one of several species that exhibit directional asymmetry at the air-sea interface. All species of Portuguese man-o-war in the genus *Physalia* exhibit directional left-right-asymmetry. Expanding these investigations to this species will allow us to compare the role of asymmetry across clades.

*Velella* are a critical component of the sea surface ecosystem, providing food and substrate for a host of free-floating and rafting animals on the high seas (Savilov 1956, Helm 2021a,b). Understanding the mechanistic and biogeographic impacts of dimorphic asymmetry in this species is important for our broader understanding of neustonic ecosystems. Beyond determining that and left- and right-handed *Velella* are differentially distributed, belonging to the same species, and that differences in asymmetric distribution are due to wind sorting, this work both elucidates a decades-old hypothesis on the consequences of *Velella* handedness and demonstrates that community and traditional science can be combined to effectively study the ocean’s surface ecosystem.

## Methods

### Biogeography using community science

Global observations of *Velella velella* were collected by iNaturalist observers and several characteristics were assessed, including sail direction, for each observation by volunteers on the crowd-source platform Zooniverse as described in Iwanicki & Helm 2025. In short, a total of 11,115 *Velella velella* observations that were available as of May 9, 2024 were imported to Zooniverse where 1,169 volunteers participated in data curation. Overall, 6,421 *Velella velella* observations containing one animal in a single image were assigned sail direction with observations covering the global extent of the species distribution. At sea observations from inside and outside of the North Pacific Subtropical Gyre were collected by professional and citizen science partners following methods described in Chong et al. 2023 across two expeditions, i) The Sea Education Association (SEA) cruise S304 aboard the SSV Robert C. Seamans and ii) the sailing crew accompanying long-distance swimmer Benoît Lecomte (https://benlecomte.com/) as he swam through the North Pacific Subtropical Gyre (The Vortex Swim). Photographs of filtered neuston net tows were analyzed in JMicroVision v1.3.5 and *Velella velella* abundance and sail direction were tabulated using the point count feature. A total of 176 *Velella velella* were observed with sail direction described across 27 stations and the results were mapped in RStudio v2024.12.1.

### DNA sequencing

Samples were collected from the coastline of North Carolina in October 2024 and Monterey Bay, California and preserved in >95% EtOH until processed. DNA was extracted from whole or dissected animals using EZNA Mollusk Kit (Omega Bio-tek) following the manufacturer’s recommendation. Whole genome sequencing was performed by Novogene Corp. using the NovaSeq X Plus Series (PE150) to a target read abundance of 20M per sample. Additional deep sequencing was performed on one specimen (NCV22) for a total of 620M PE150 reads. Raw reads were quality assessed using FastQC v.0.12.0 and trimmed to remove Illumina adapters using Trimmomatic, v. 0.39 (Bolger et al. 2014).

### Phylogenetics

Mitochondrial genomes were assembled for each individual, including *Porpita porpita* as an outgroup (Yale Peabody Museum Catalog number: [YPM_IZ_111244](https://collections.peabody.yale.edu/search/Record/YPM-IZ-111244)) using GetOrganelle v1.7.7.1 (Jin et al. 2020) and aligned using mafft v.7.525 (Katoh & Standley 2014). A maximum likelihood tree was inferred using IQ-TREE v2.4.0 with ultrafast bootstrapping. The best fit model TVM+F+I+R3 was automatically selected for the mitochondrial genome alignment.

### Population genetics

Population genetic analyses were performed using trimmed reads mapped to a tree-informed deduplicated transcriptome following the methodology detailed in Church et al. 2025. In short, a transcriptome was assembled from publicly available *Velella velella* reads (SRR8115525) using trinity v.2.8.5 with default settings and deduplicated using the treeinform pipeline (Guang et al. 2021, Munro et al. 2022). Trimmed genomic reads were mapped to the transcriptome using bwa v.0.7.18 and sorted with picard v3.4.0, mapped reads were filtered and merged using bcftools v1.20, indexed using tabix v.1.22, and population genetic statistics were inferred using pixy v.2.0.0. Single nucleotide polymorphism (SNP) weighted average *Fst* was calculated and Kruskal-Wallis test with a *post hoc* Dunn’s test with Holm correction was performed to assess statistical significance in RStudio v2024.12.1.

### Genome size estimation

We estimated the genome size for one specimen (NCV22) sequenced to >20x coverage (620 million PE150 reads), estimated against a genome size of 3 Gb. Genome size, repeat content, and heterozygosity were estimated using GenomeScope, v. 2.0 (Ranallo-Benavidez et al. 2020) and jellyfish, v. 2.2.3 (Marçais et al. 2011). GenomeScope was run on the combined set of R1 and R2 reads, and trimmed as described above.

### Velella drift in the ocean

We are not aware of any *Velella* tracking data in the open ocean, so we used results of the Mackie (1962) experiment to derive how *Velella* responds to the local wind. Our experience with marine debris (Maximenko and Hafner, 2024) suggests that under a broad range of parameters, drift speed of an object floating on the ocean surface is a combination of the surface current and additional slip relative to the water caused by the direct wind force. This slip is often proportional to the wind speed (e.g., Lee and Maximenko 2025).

By converting information of Mackie (1962) into a practical coefficient, we derived that a left-handed *Velella* were drifting at a 37 degrees angle to the left from the wind direction at a speed of 3.5% of the wind speed. We assumed that for righties, the response would be the same but the drift velocity vector would be to the right from the wind direction (Fig. 1A). This estimate may lack accuracy because the experiment was conducted in a tidal pool but it helps to demonstrate effects of the handedness and interpret observational data. In more comprehensive consideration, the coefficient may depend on the sea state and size of the individuals. The coefficient was then used to derive mean *Velella* streamlines (Fig. 1C,D) using ocean wind and currents averaged over a long period of time.

For the analysis based on reports submitted through Zooniverse and presented in Fig. 4, we used only reports describing single *Velella* observations. Although most of *Velella* strandings were happening in groups of various sizes, characterization of fractions with different orientations was difficult.

## Supporting information

Supp File A: zooniverse data

Supp File B: gyre images and count files

Supp File C: mitochondrial ML tree

## Acknowledgements

We would like to thank the numerous iNaturalist observers and identifiers, Zooniverse volunteers, with special thanks to Larry Jensen, Zane Warden, Ashleigh Swett, and Amanda Cogan Barber for feedback; the Zooniverse support team; Deborah Goodman with the Sea Education Association and crew; Benoît Lecomte and the Vortex Swim crew; Steven Haddock at Monterey Bay Aquarium Research Institute; Ary Puentes with GO-SEA. This study is indirectly supported by the NASA GO-SEA project (www.goseascience.org) through grant 80NSSC21K0857.

## Author Contributions

R.R.H. conceived of the study. T.I. and R.R.H. and N.M. were involved in concept & design, and drafting of the original manuscript; T.I. developed the Zooniverse project and performed molecular analyses; N.M. conducted the oceanographic analyses; S.H.C. and C.W.D. developed software and were involved in the design of molecular analyses; all authors contributed to drafting/revisions and approval of the submitted manuscript.

## Supplementary

Supp. File A: csv containing all singleton observations that include all meta-data, including count, condition, sail direction, url to original iNat, whether the observation was manually corrected, etc.

Supp. File B: images and corresponding jmv files for *Velella* counts from neuston net tows collected during i) The Vortex Swim Expedition and ii) the Sea Education Association cruise S304.

Supp. File C: tree file for aligned mitochondrial genomes of *Velella* collected in Monterey Bay, California (MBV) and North Carolina (NCV) with *Porpita porpita* outgroup.

**Table 1:** Contingency table of Pearson’s Chi-squared test of Zooniverse determined data of *V. velella* handedness from global sampling efforts on iNaturalist. Left-handed animals are ∼5.5 more abundant in our dataset than right-handed animals, thus we expect this same ratio in the northern and southern hemispheres.

|  | North (obs) | North (exp) | South (obs) | South (exp) |
| --- | --- | --- | --- | --- |
| Left | 5000 | 4610 | 443 | 833 |
| Right | 438 | 828 | 540 | 150 |

**Fig.**
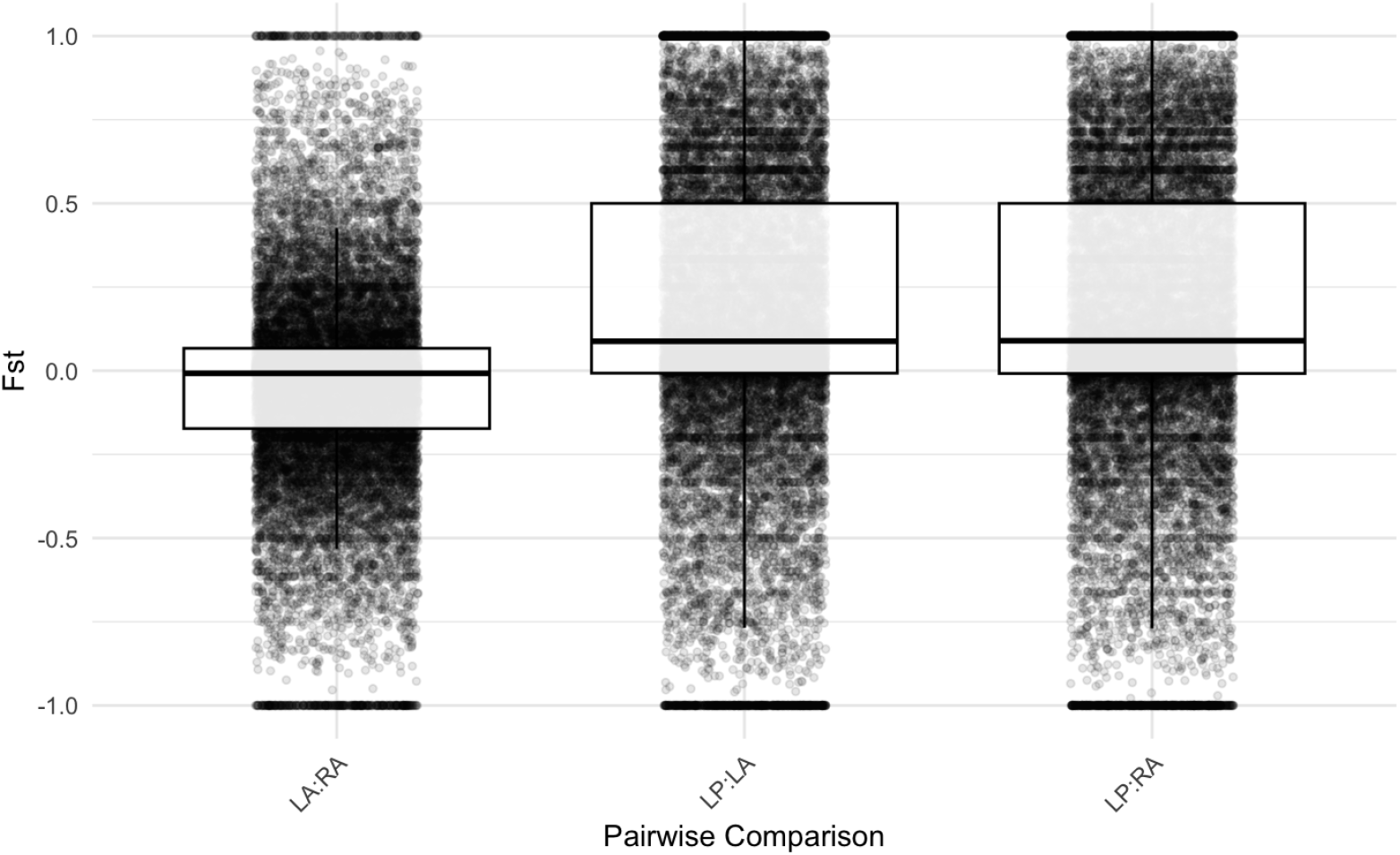
Pairwise box-plot of *Fst* values for *Velella velella* that are either left-handed (LA) and right-handed (RA) collected from the Atlantic Ocean and left-handed (LP) collected from the Pacific Ocean. Quality-filtered genomics reads were mapped to a tree-informed reference transcriptome assembled from publicly available data (SRR8115525).

