## Supplementary figures and images for "Global distribution patterns of an ocean-surface dwelling animal are associated with organismal mirror asymmetry"

### HelmLab_S304-009-NT_07012022_Pristine.tif

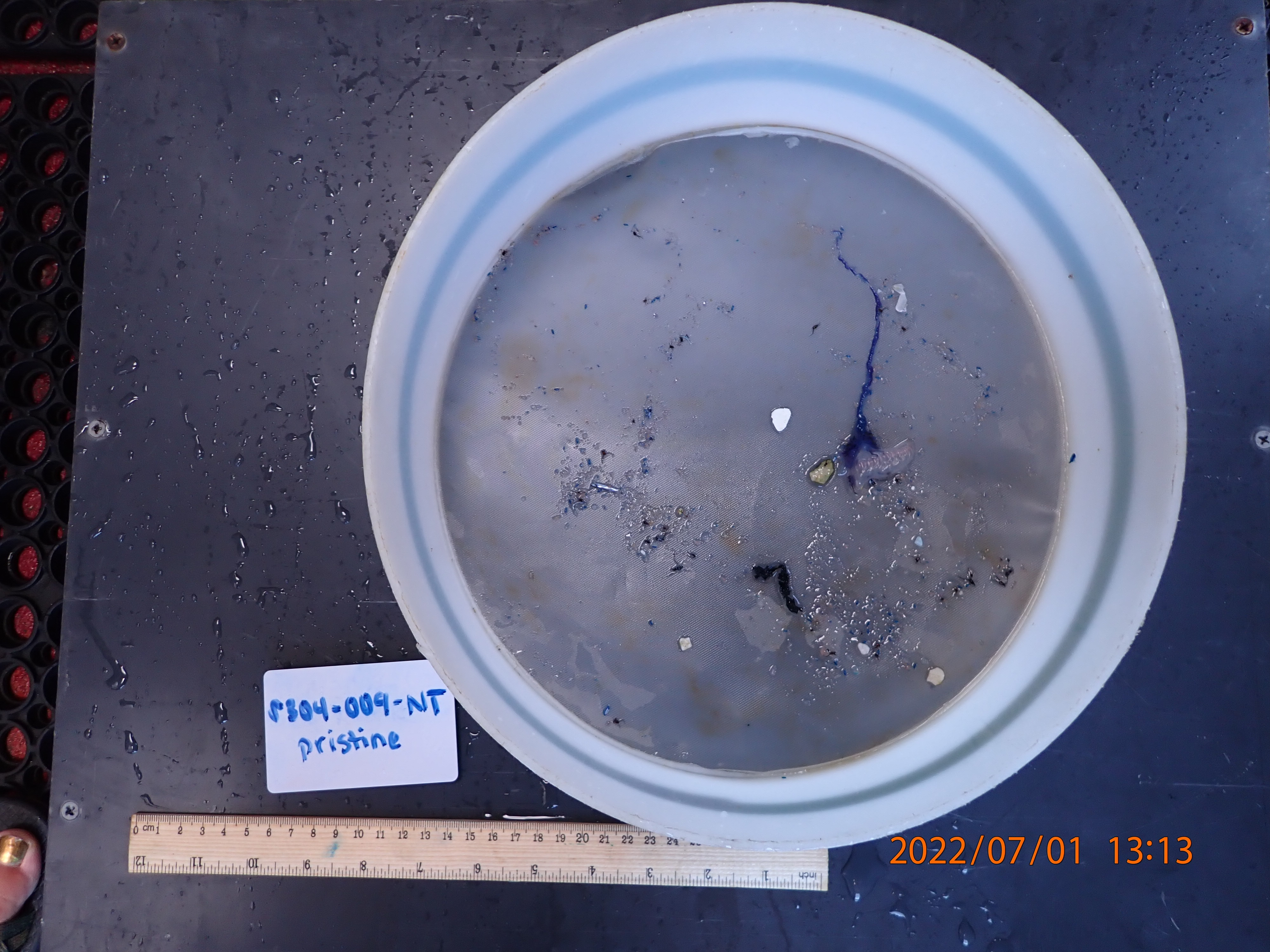

### HelmLab_S304-009-NT_07012022_Rinse.tif

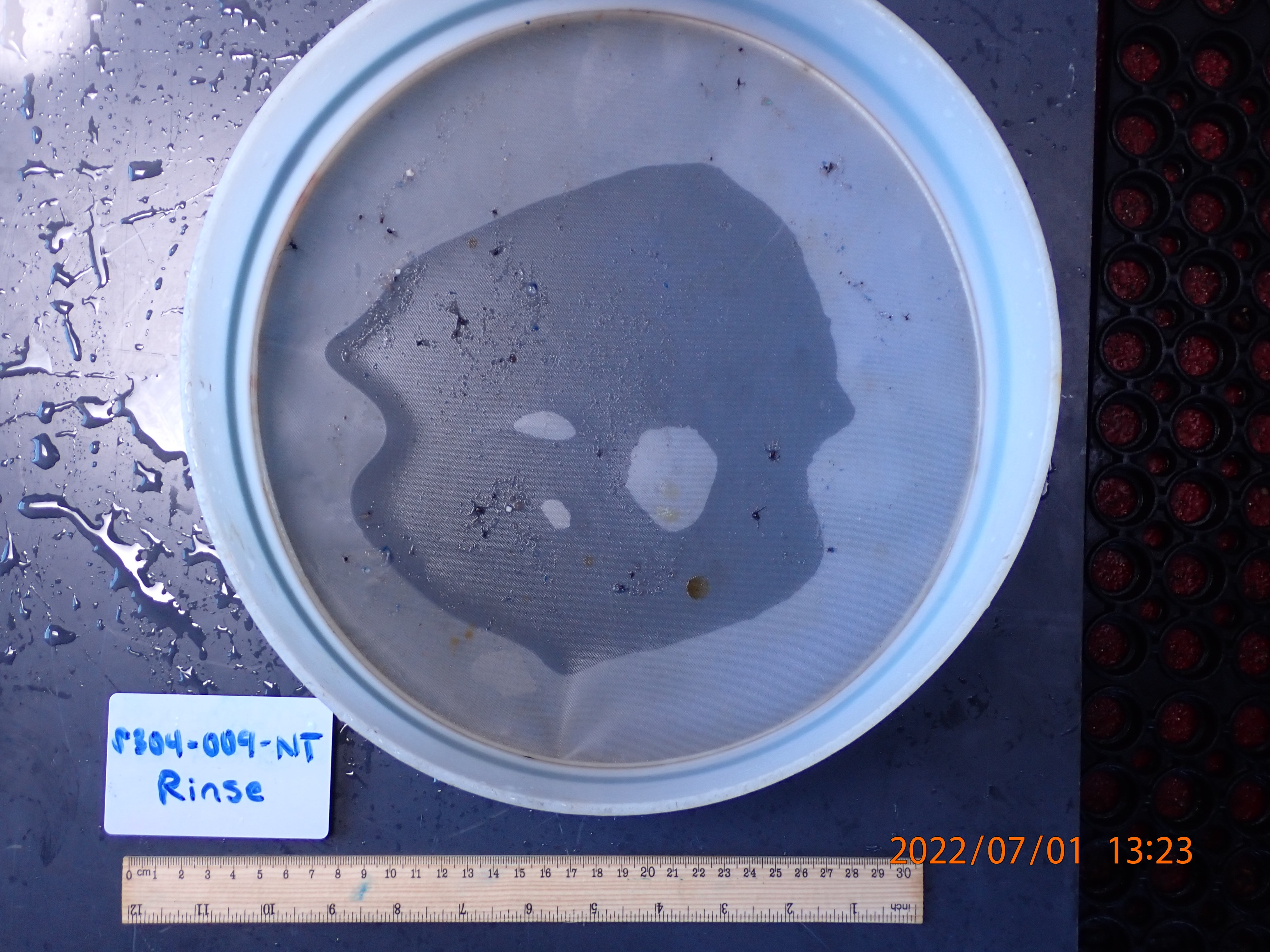

### HelmLab_S304-010-NT_07012022_Rinse.tif

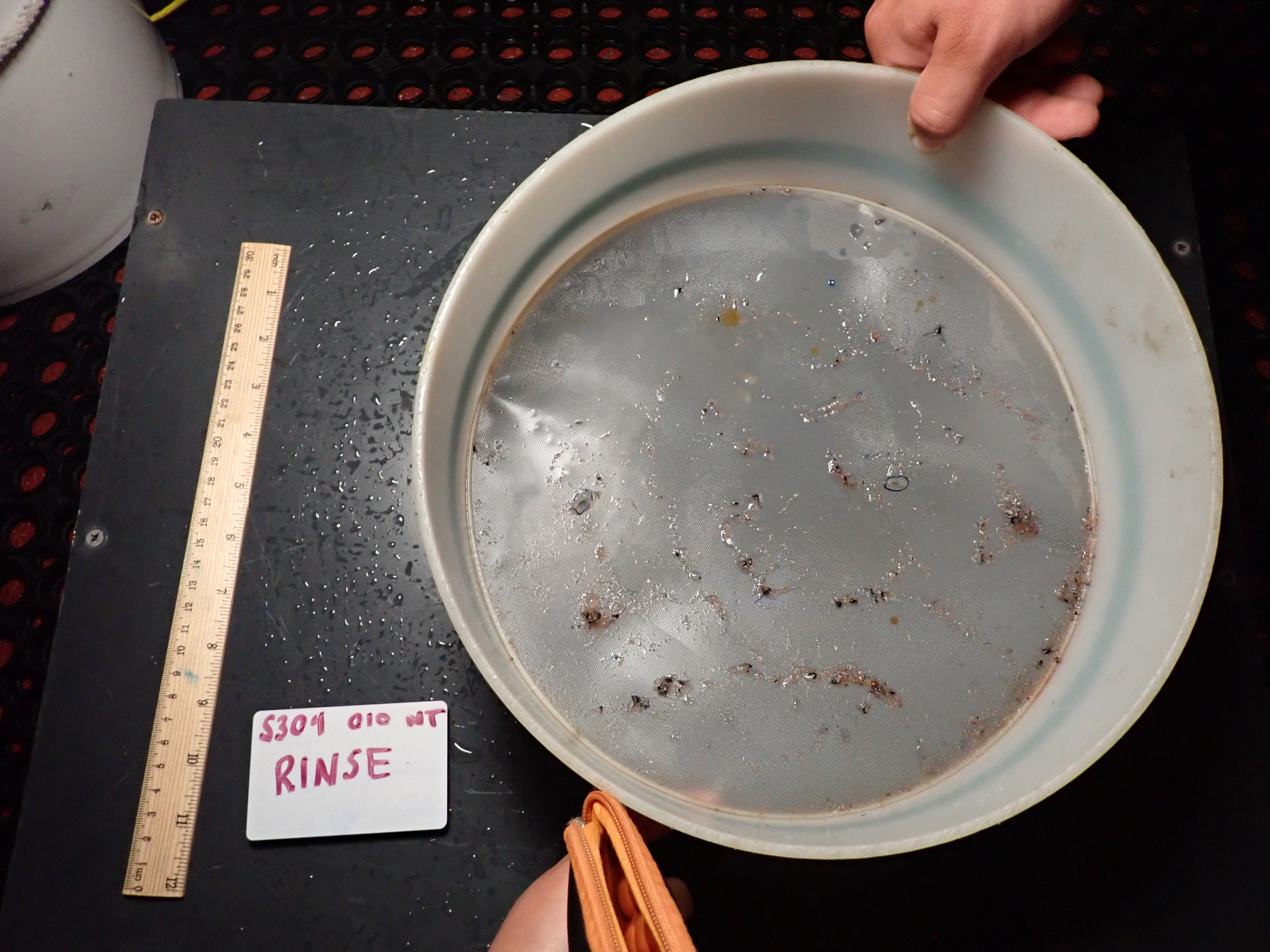

### HelmLab_S304-011-NT_07022022_Pristine.tif

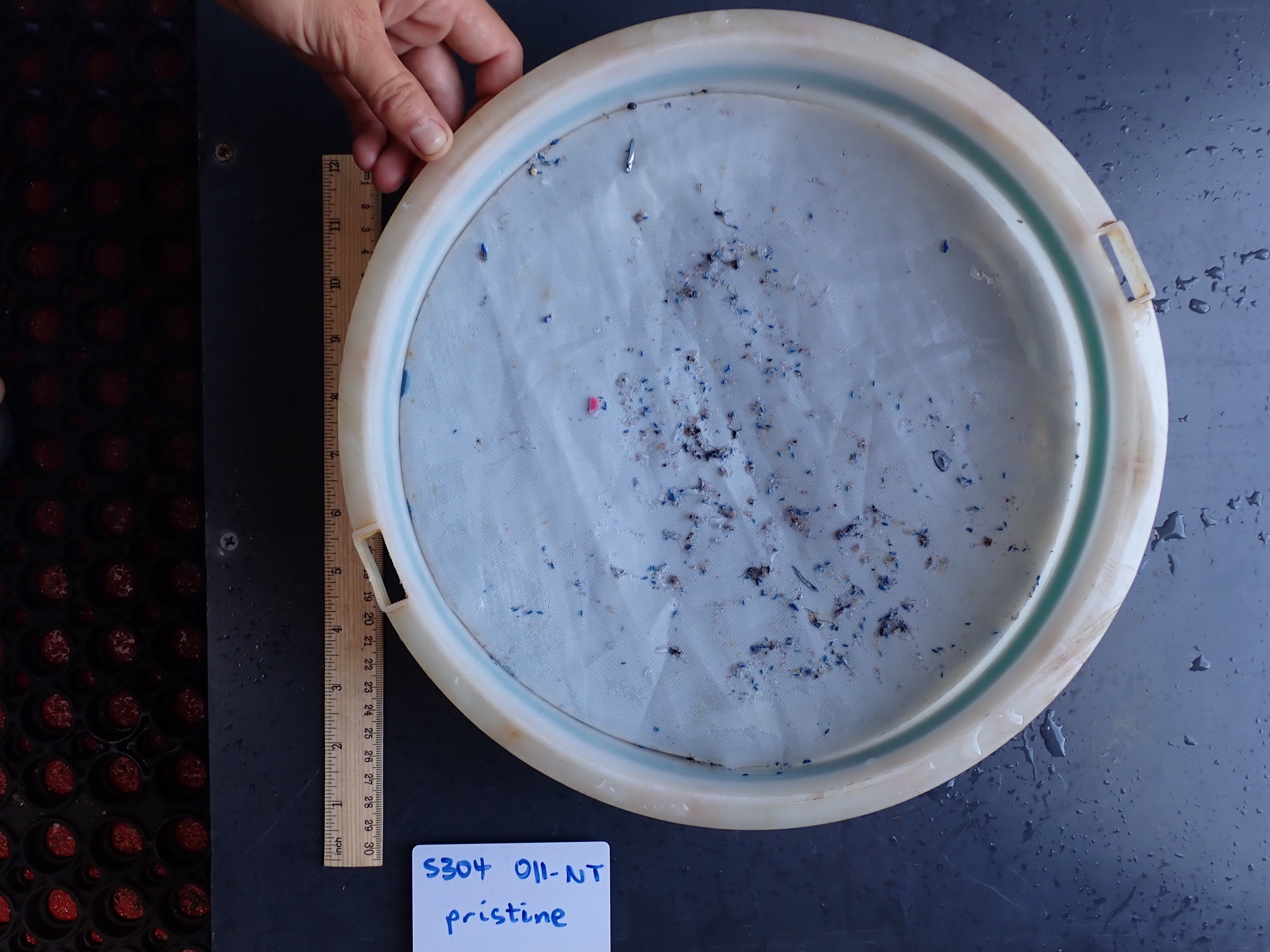

### HelmLab_S304-013-NT_07032022_Pristine.tif

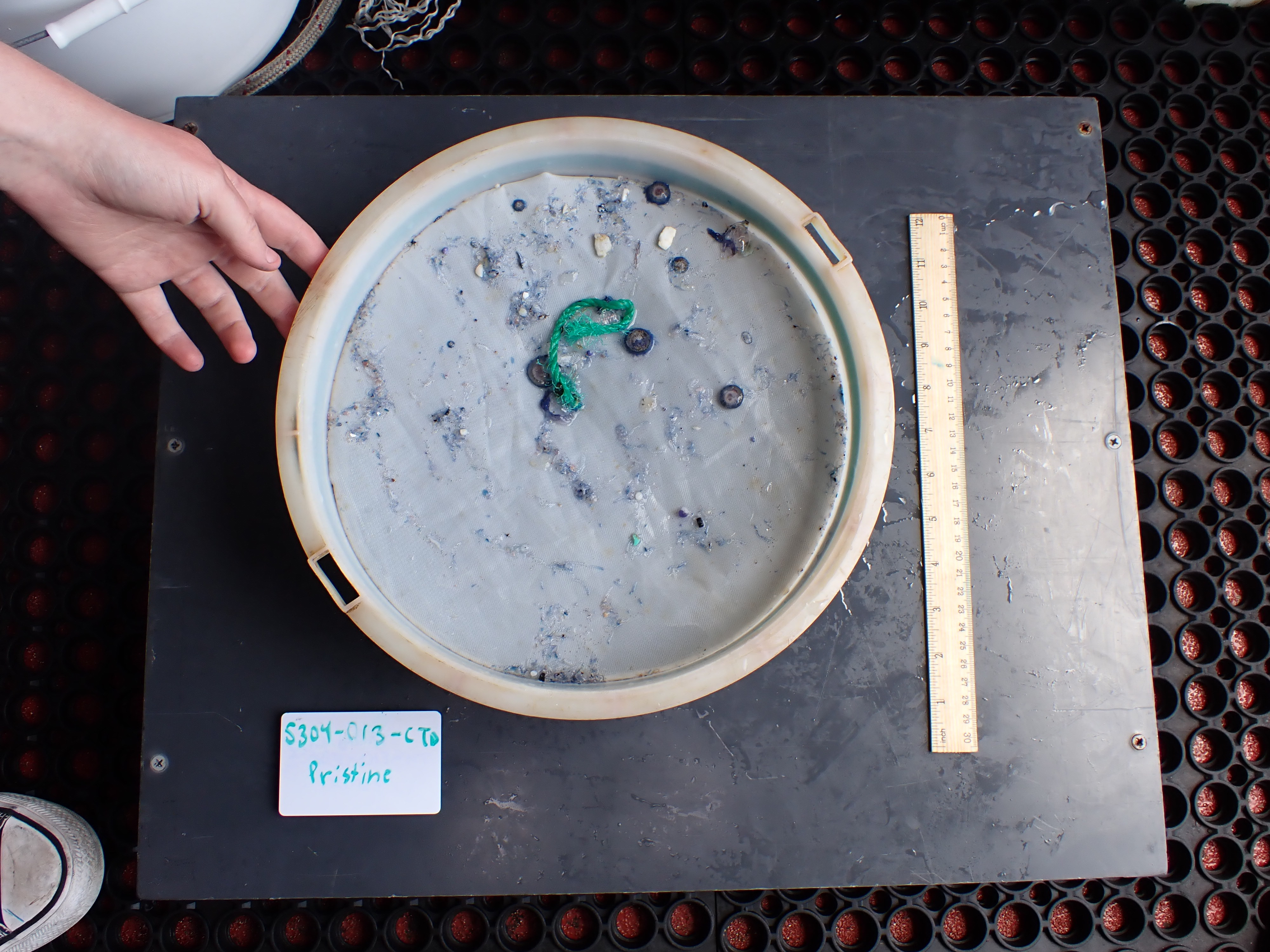

### HelmLab_S304-014-NT_07032022_Rinse.tif

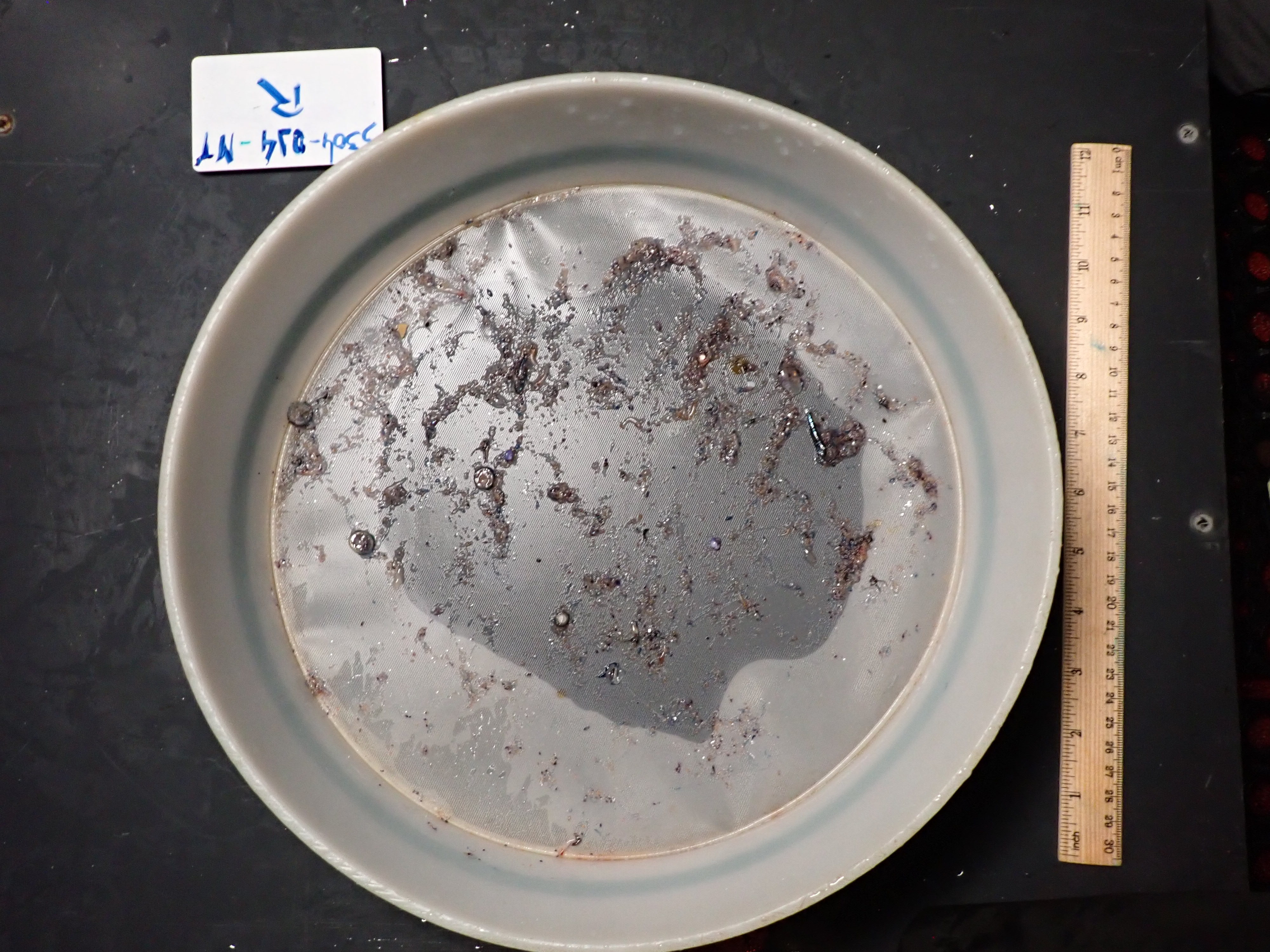

### HelmLab_S304-015-NT_07042022_Pristine.tif

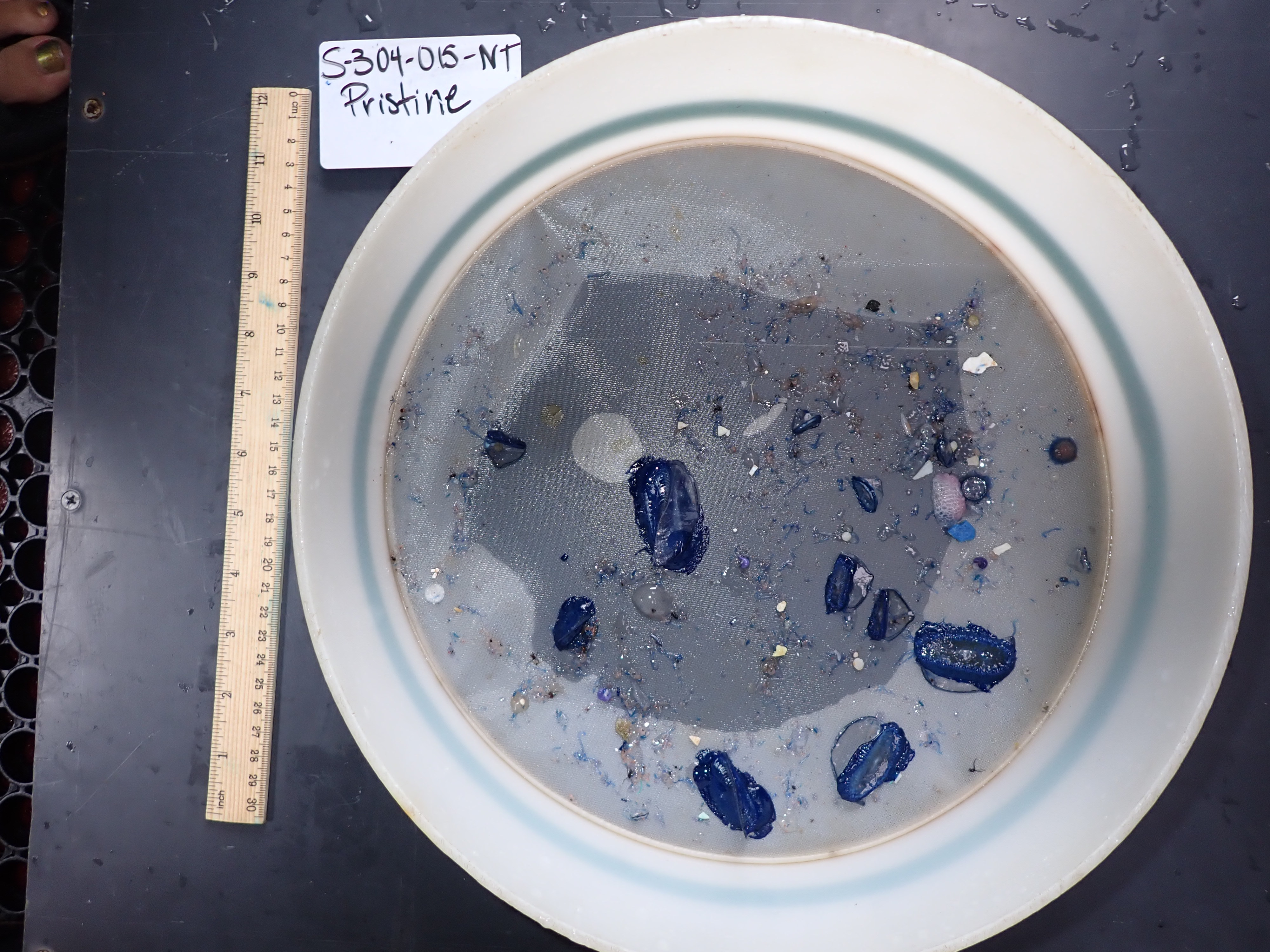

### HelmLab_S304-015-NT_07042022_Rinse.tif

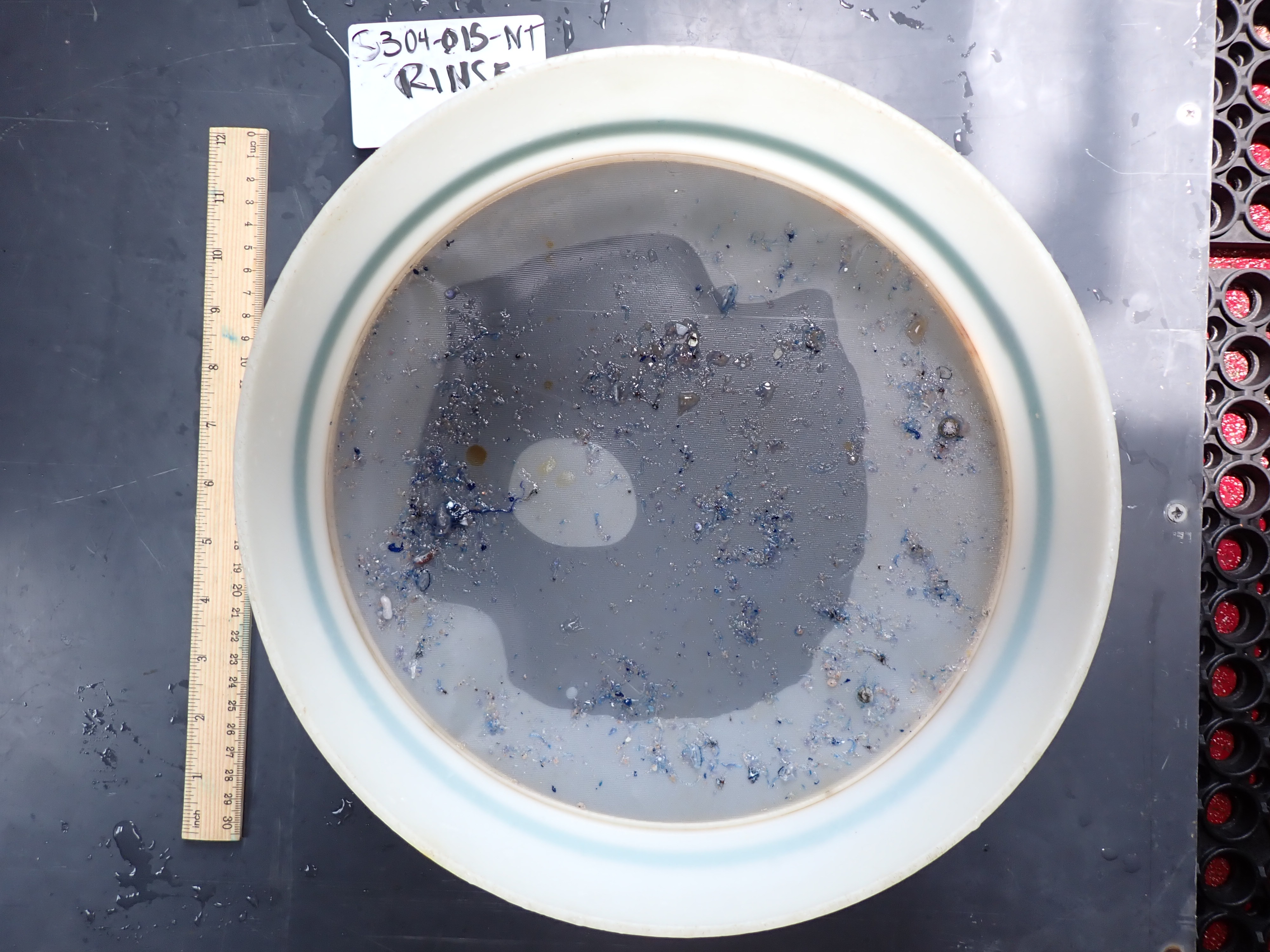

### HelmLab_S304-016-NT_07052022_Pristine.tif

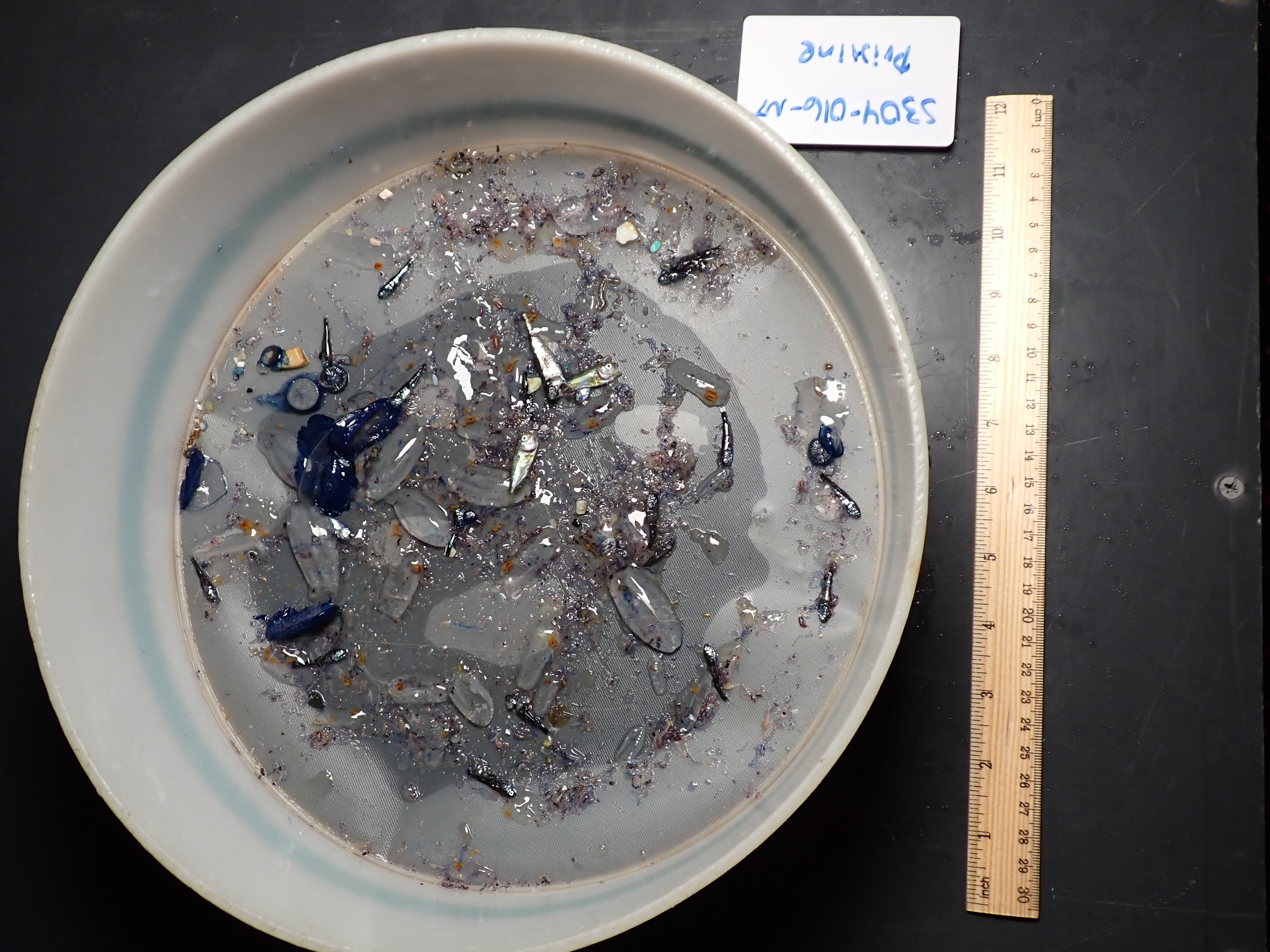

### HelmLab_S304-016-NT_07052022_Rinse.tif

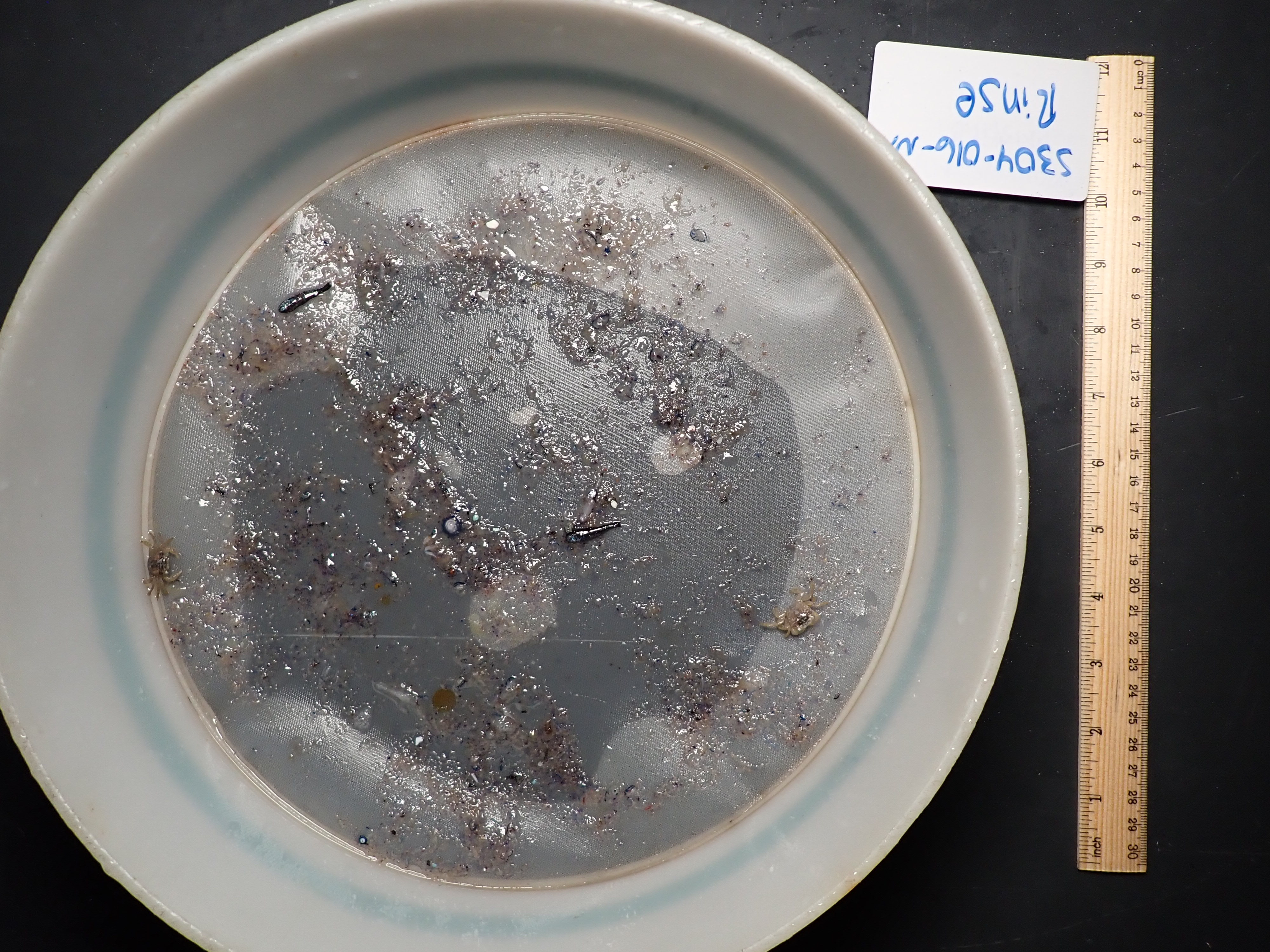

### HelmLab_S304-017-NT_07062022_Pristine.tif

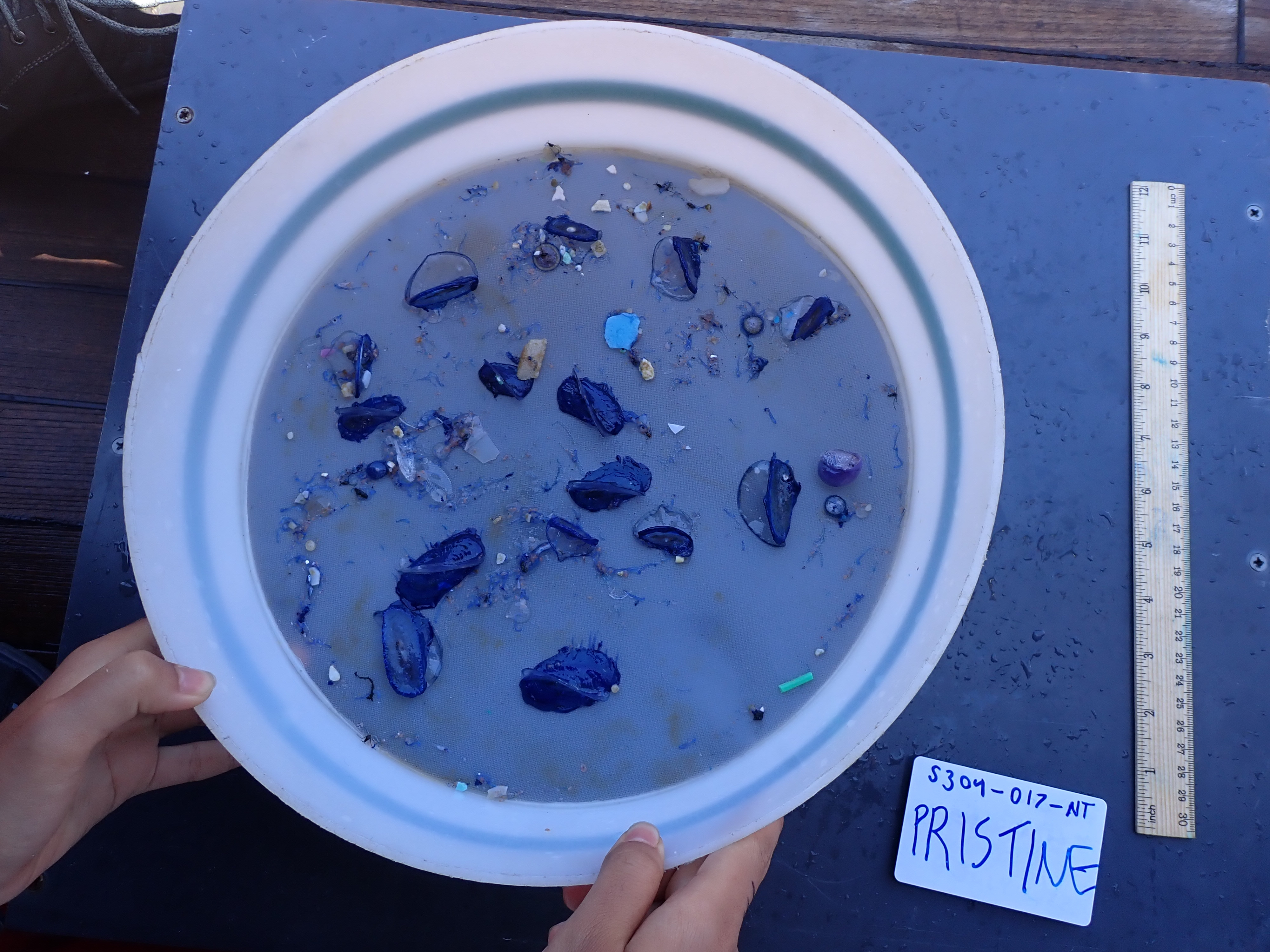

### HelmLab_S304-019-NT_07072022_Pristine.tif

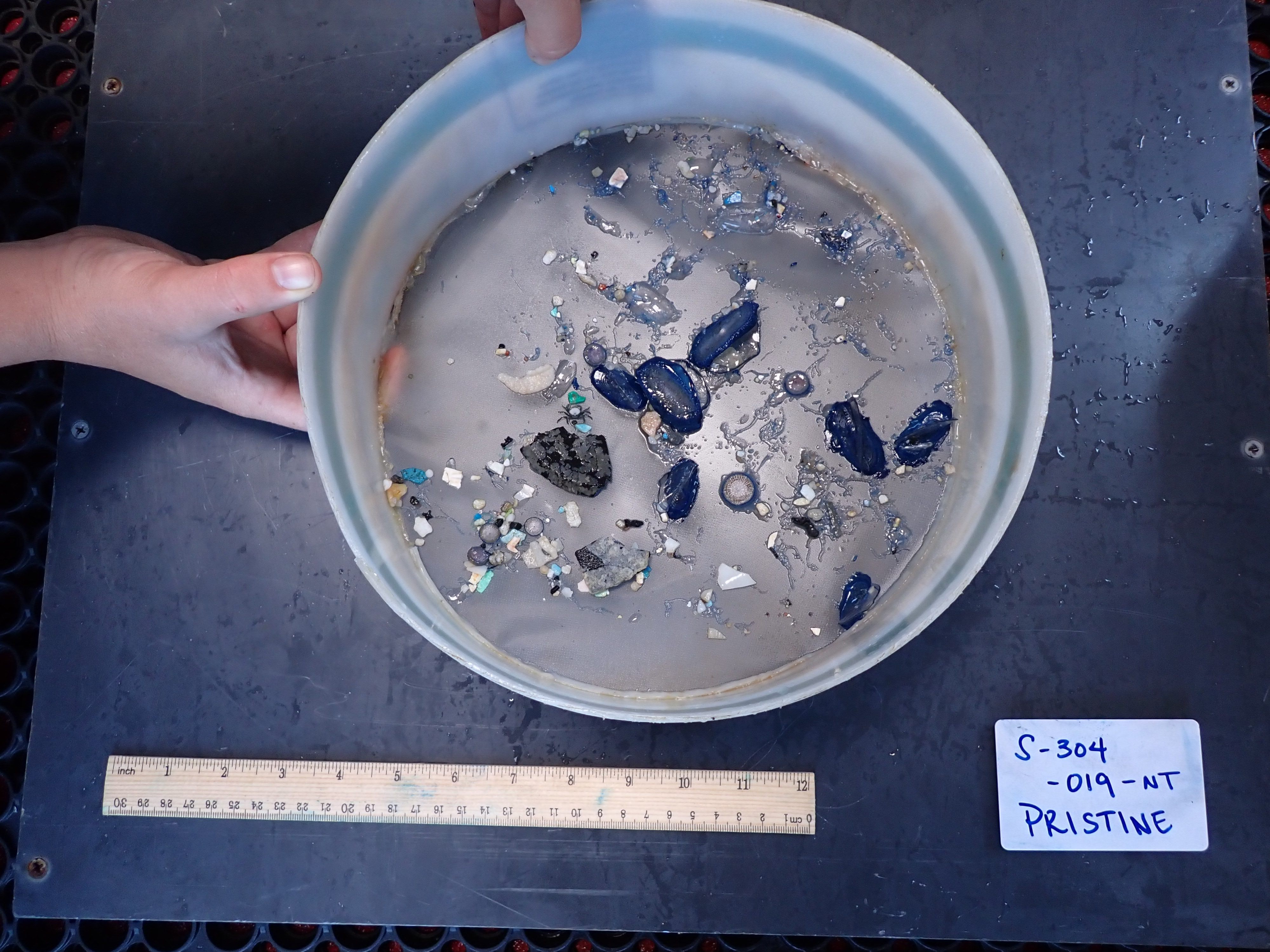

### HelmLab_S304-021-NT_07072022_Pristine.tif

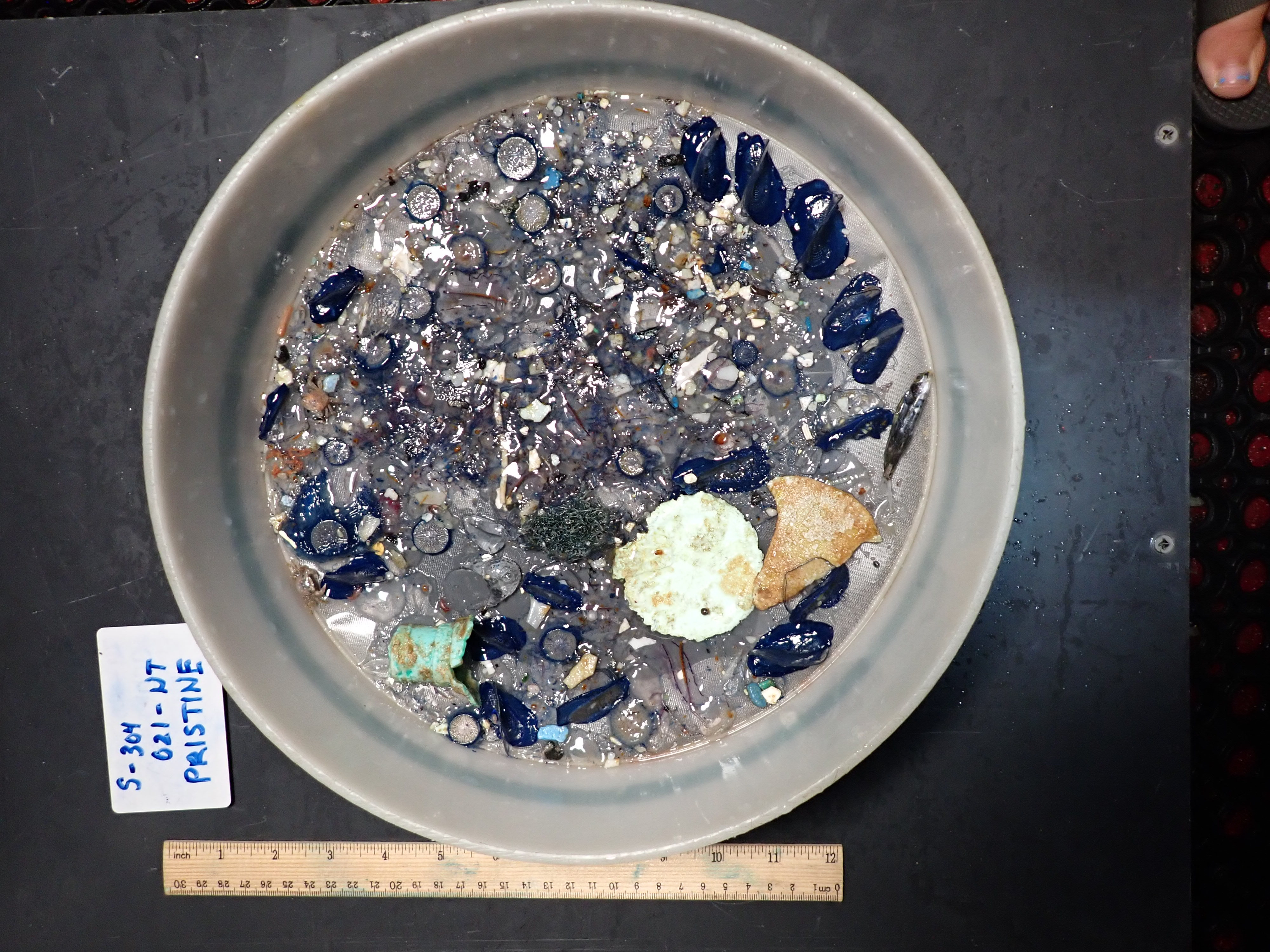

### HelmLab_S304-022-NT_07082022_Pristine.tif

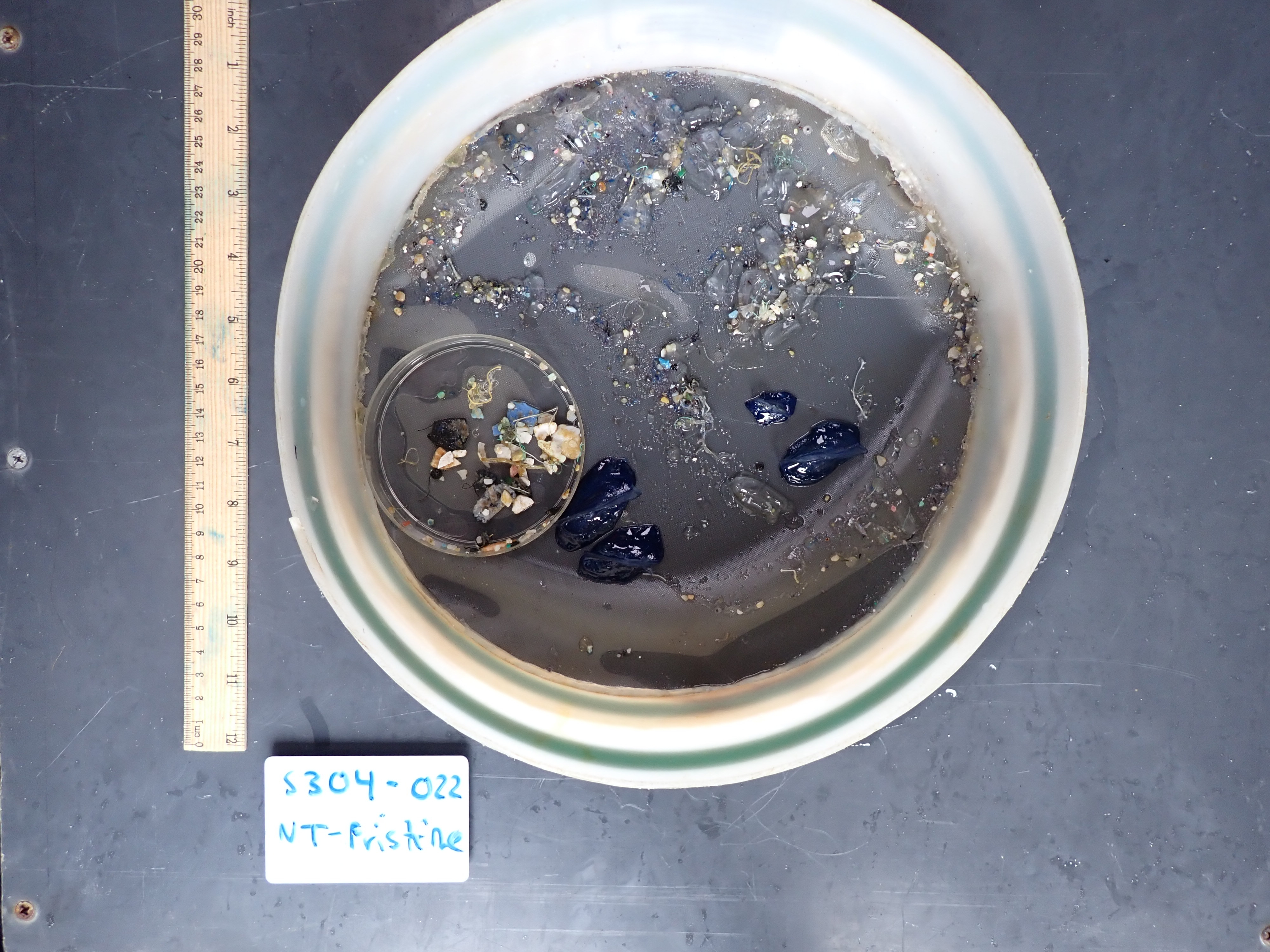

### HelmLab_S304-023-NT_07082022_Pristine.tif

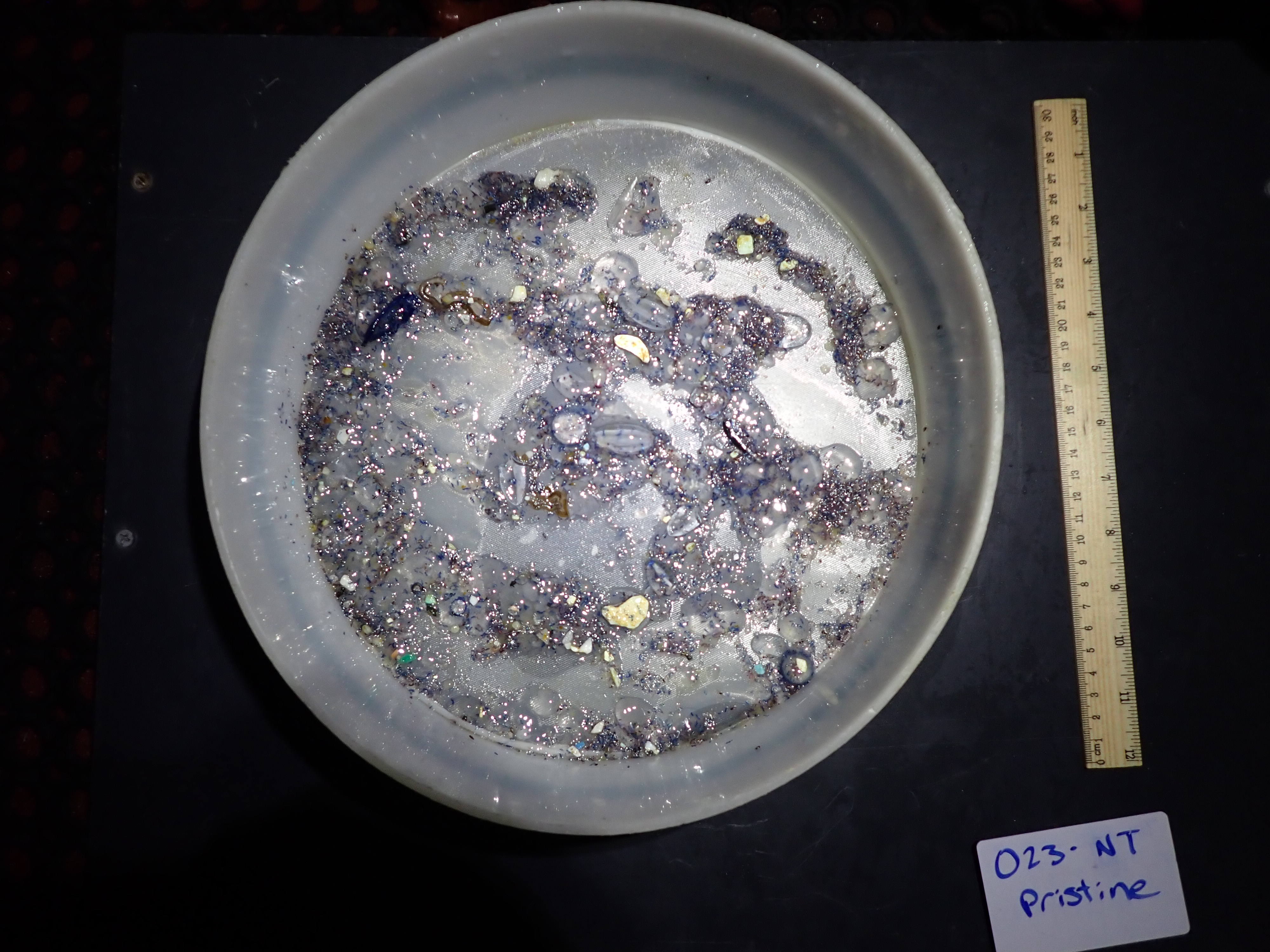

### HelmLab_S304-029-NT_07102022_Pristine.tif

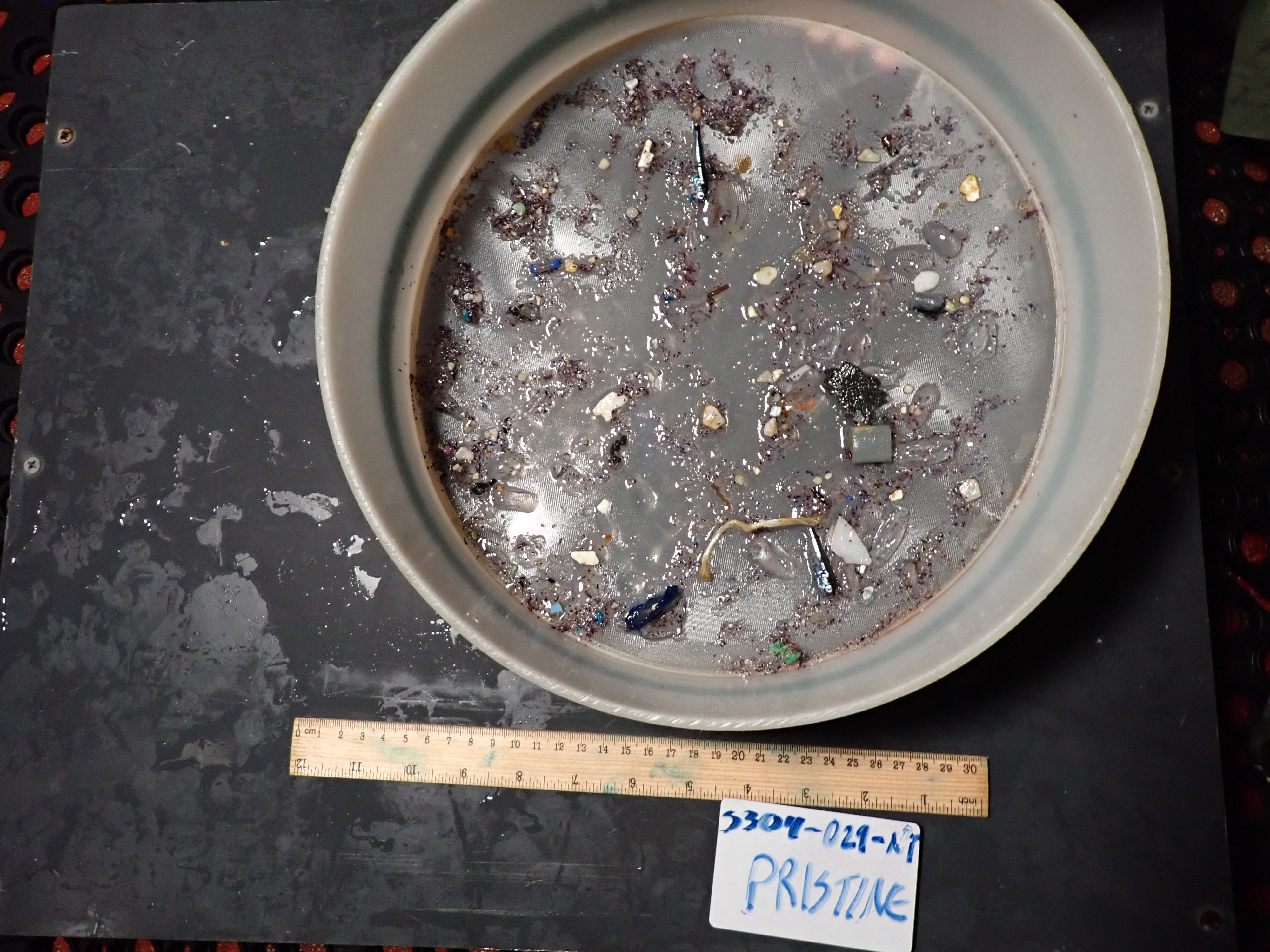

### HelmLab_S304-032-NT_07122022_Pristine.tif

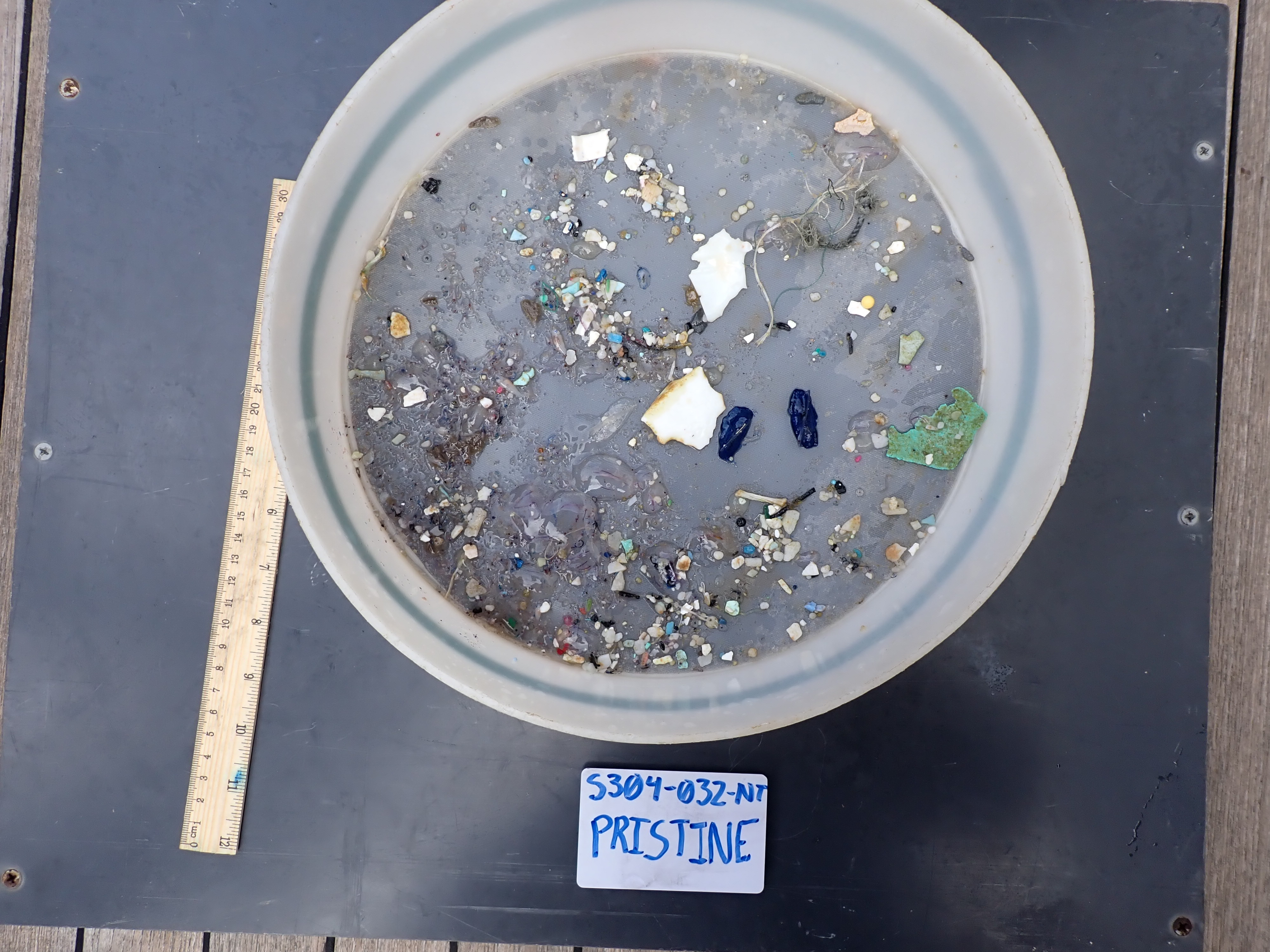

### HelmLab_S304-042-NT_07162022_Pristine.tif

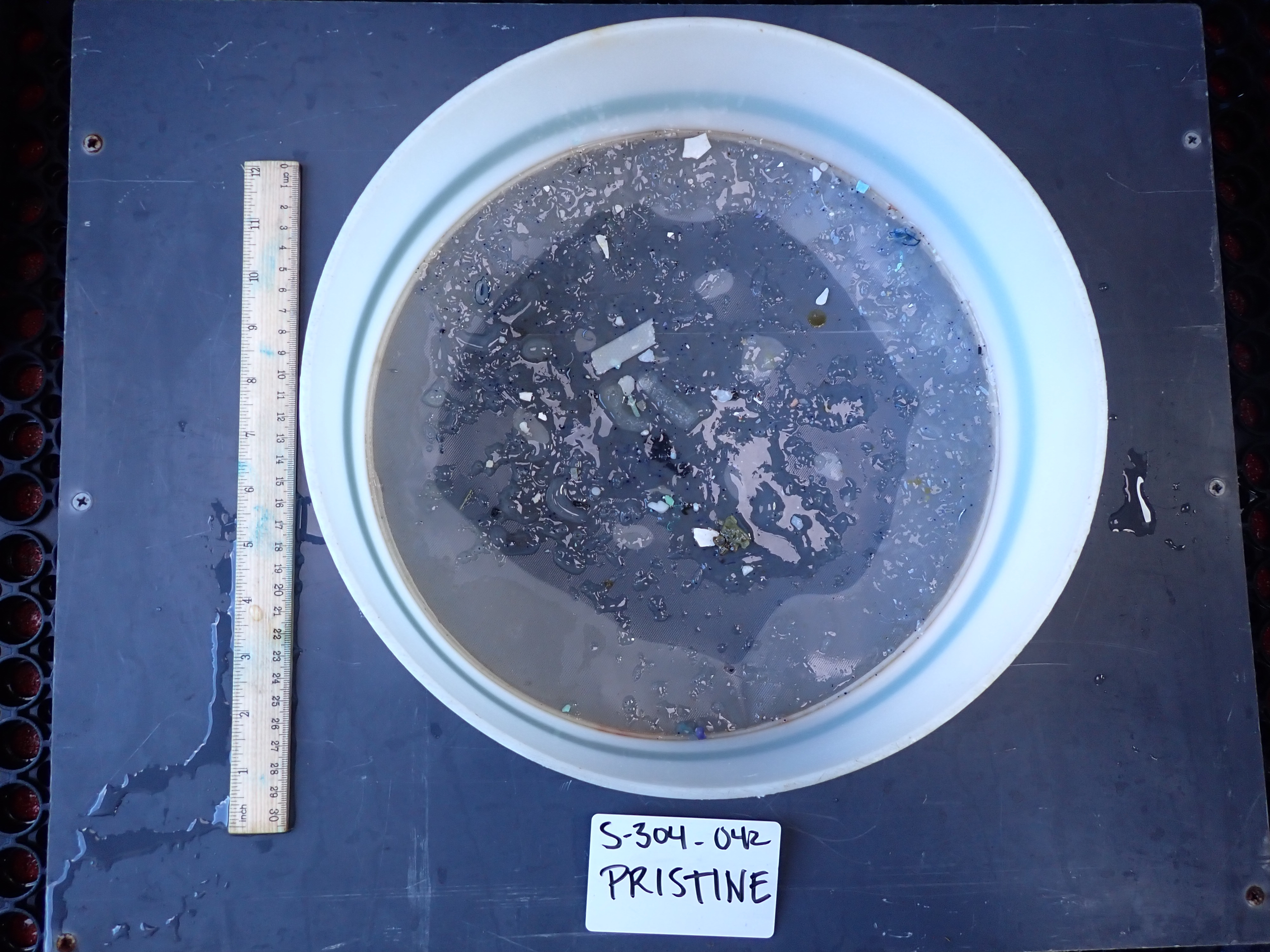

### HelmLab_S304-043-NT_07162022_Rinse.tif

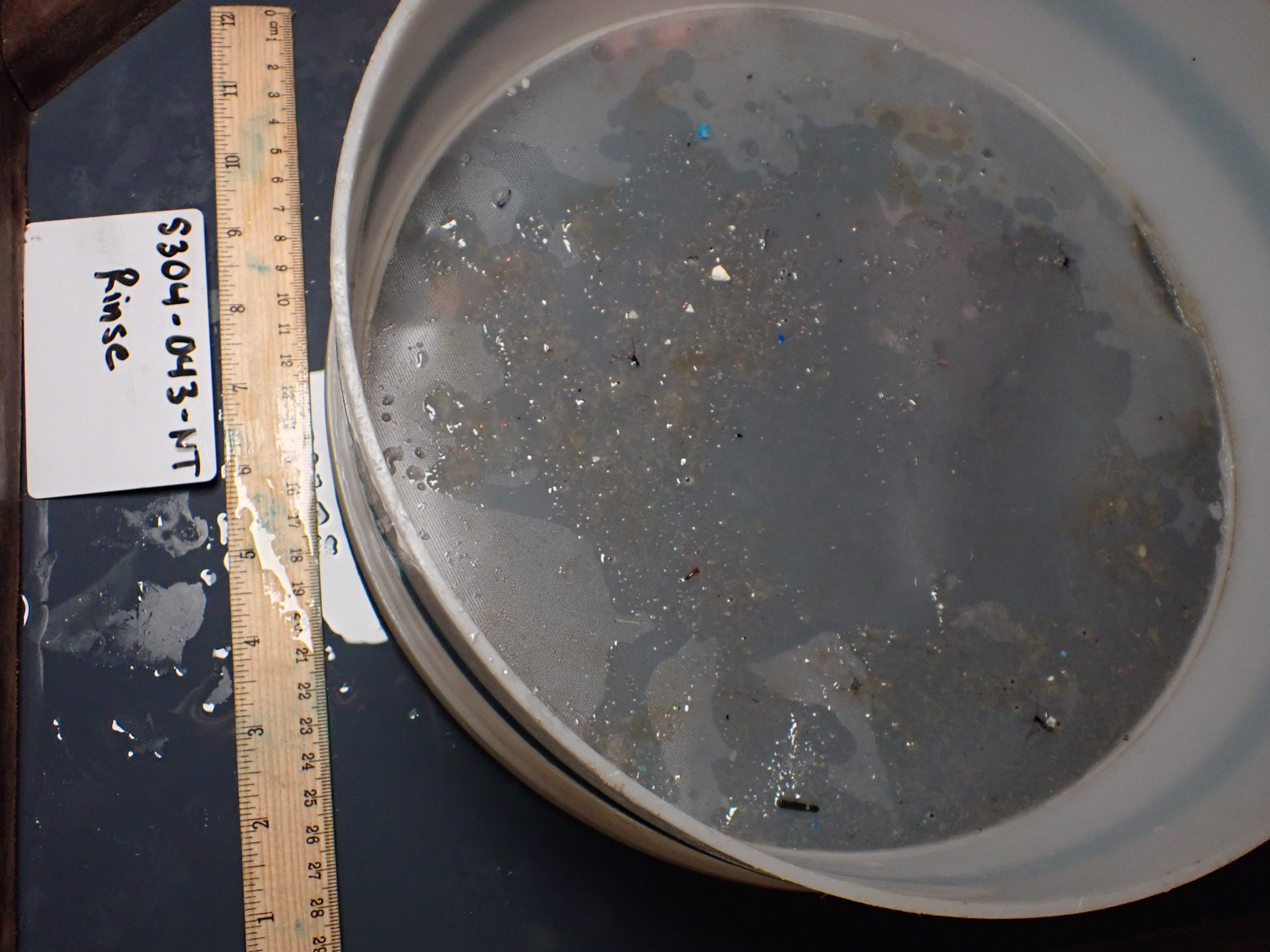

### HelmLab_S304-044-NT_07172022_Pristine.tif

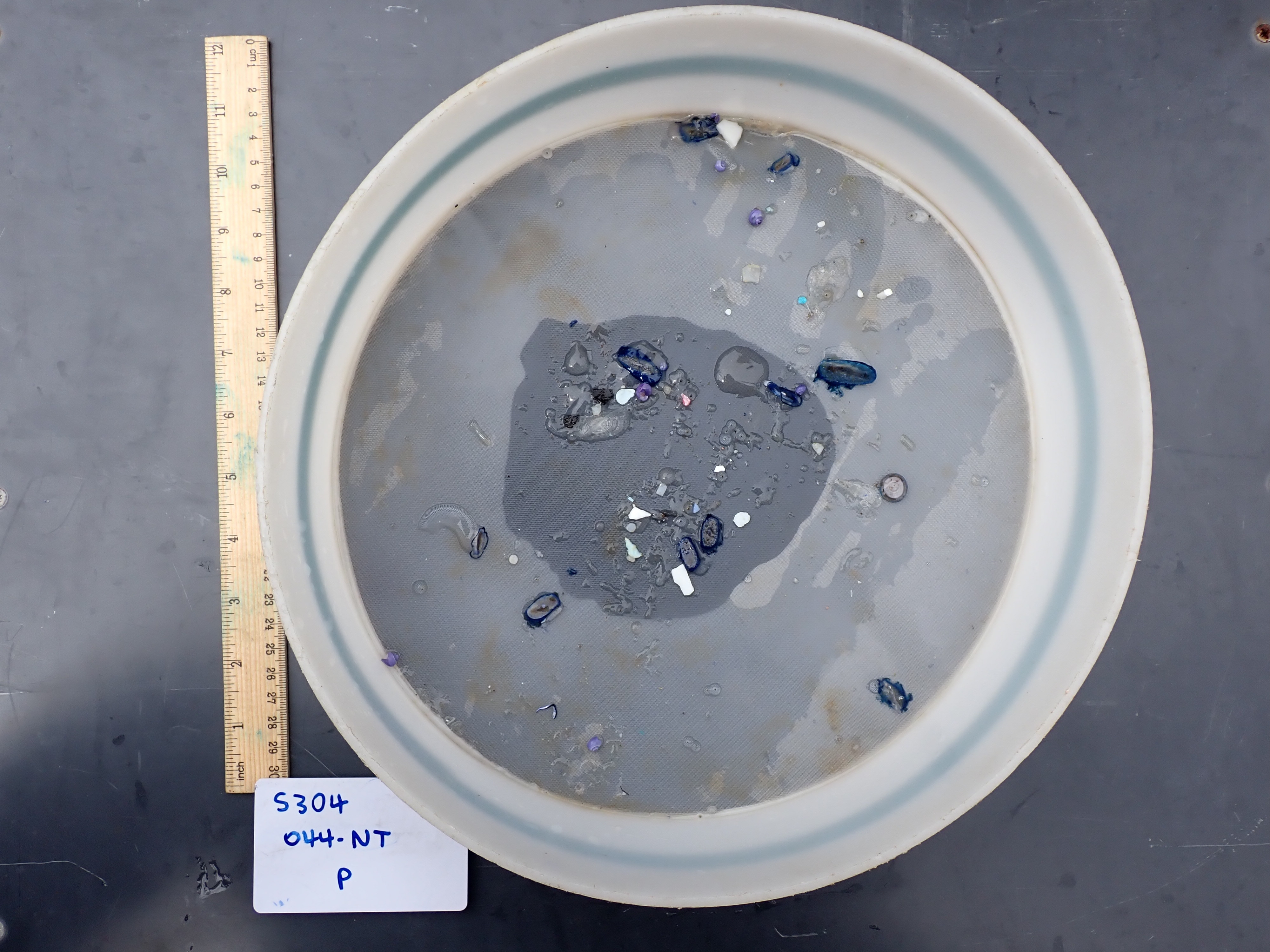

### HelmLab_S304-045-NT_07172022_Pristine.tif

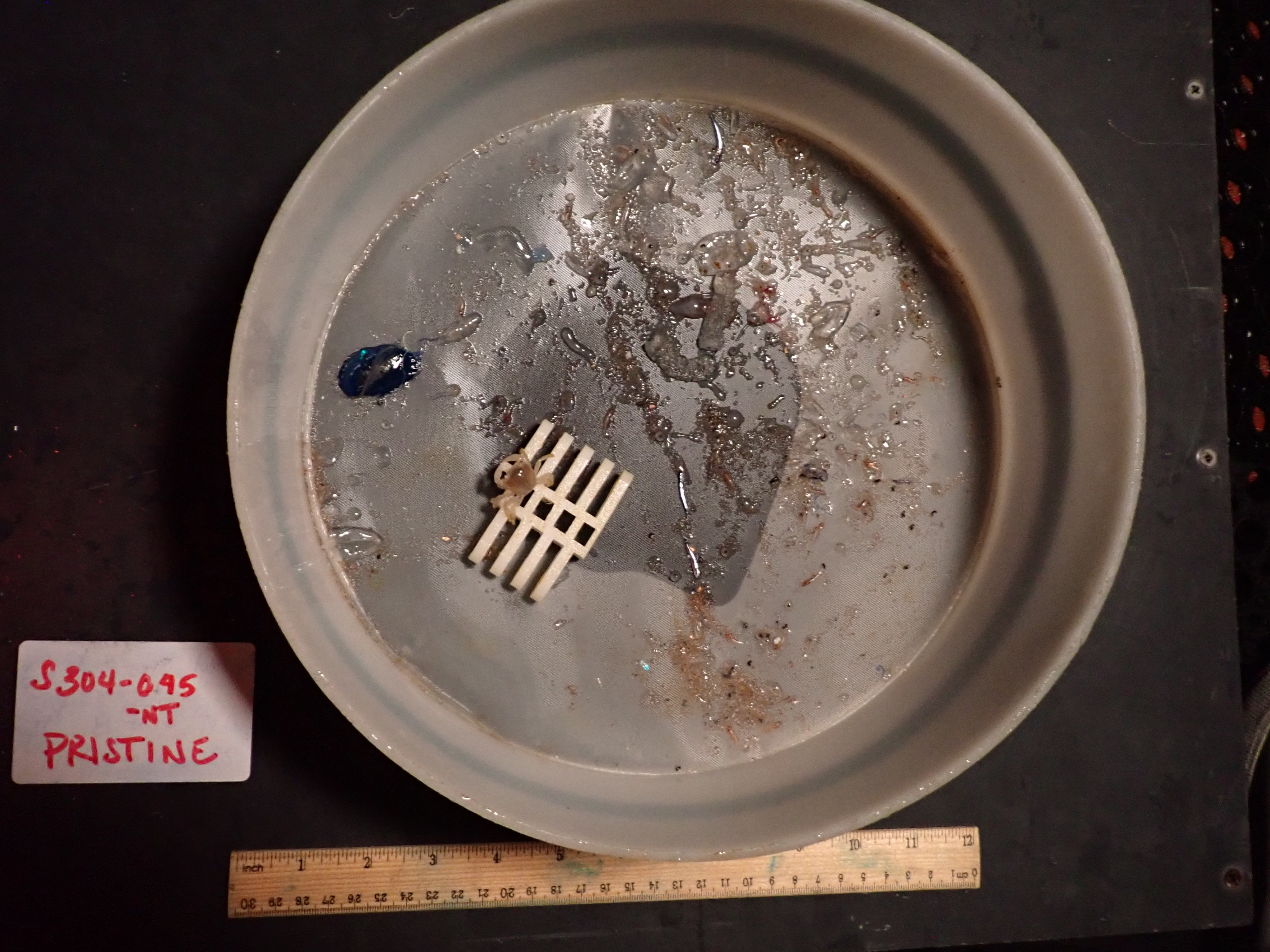

### HelmLab_S304-046-NT_07182022_Pristine.tif

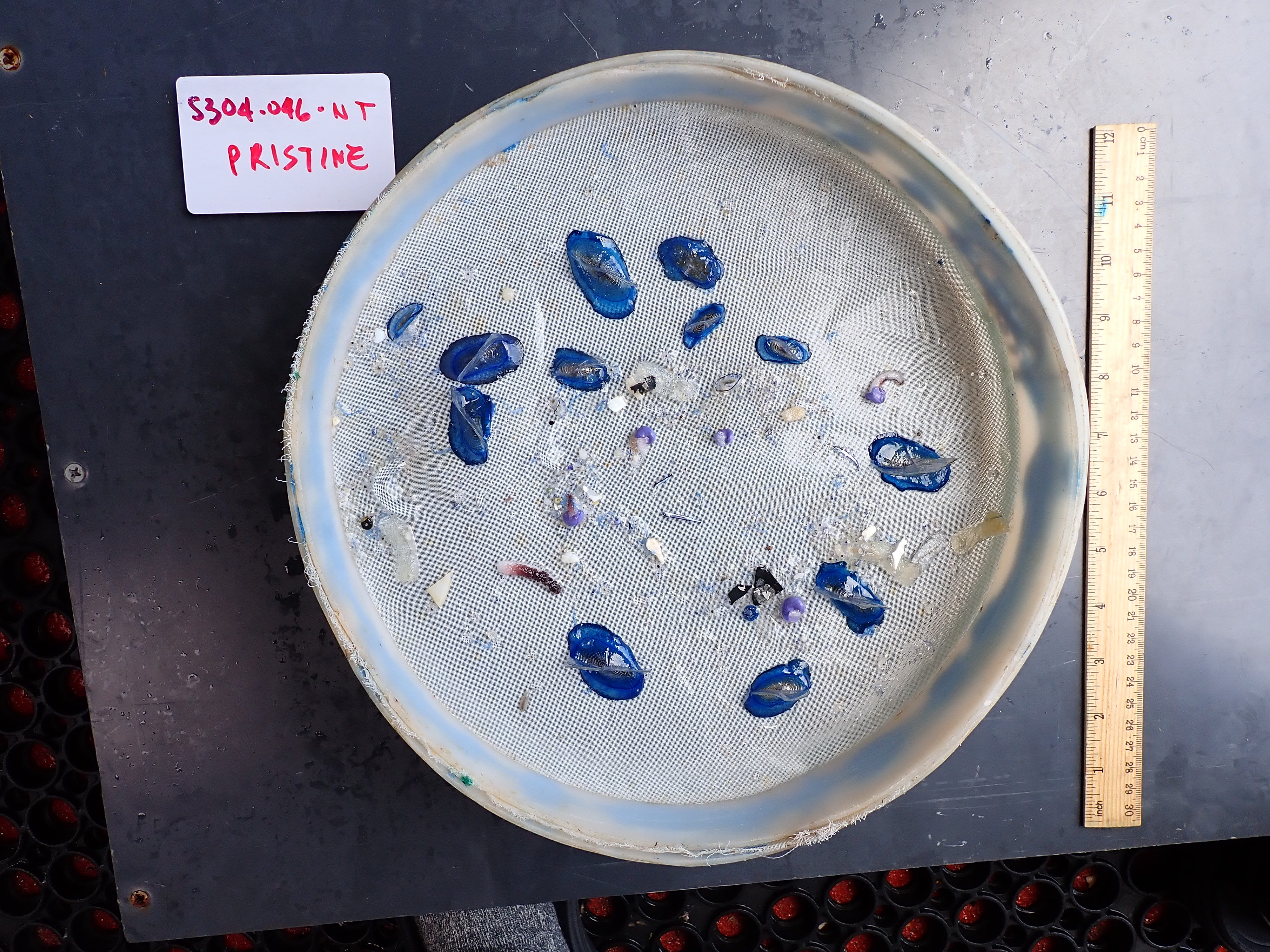

### HelmLab_S304-047-NT_07182022_Pristine.tif

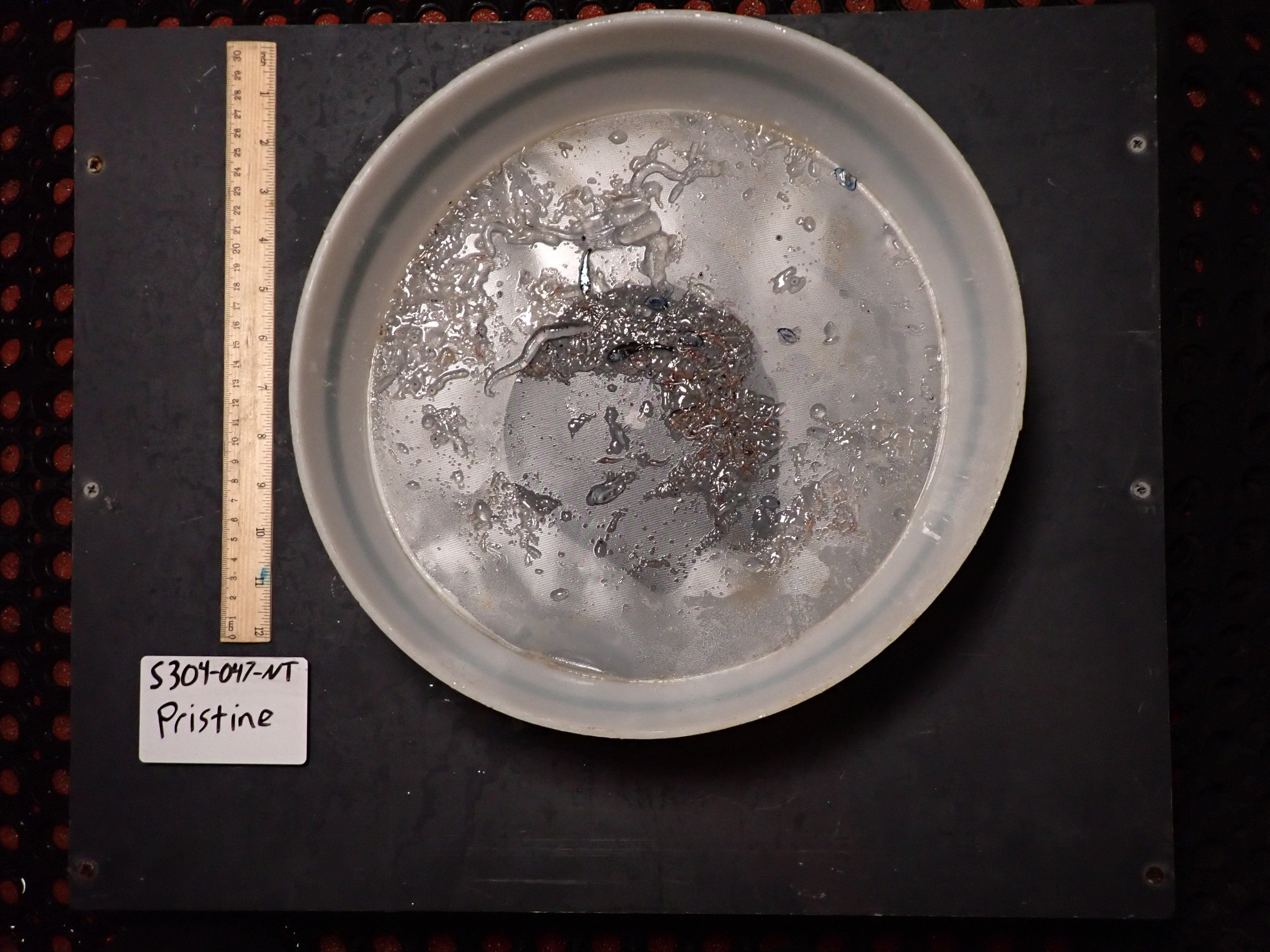

### HelmLab_S304-048-NT_07192022_Rinse.tif

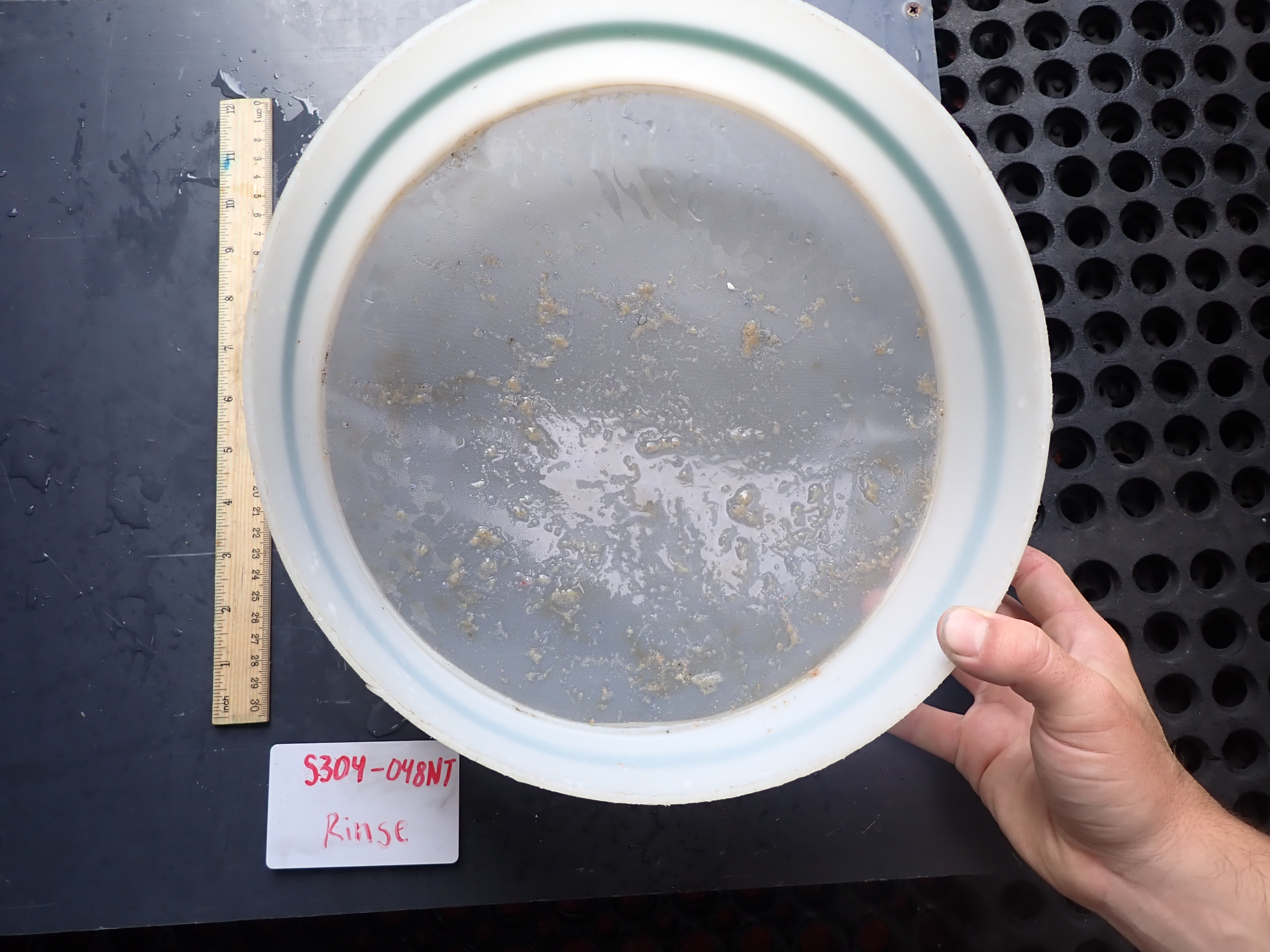

### HelmLab_S304-050-NT_07202022_Pristine.tif

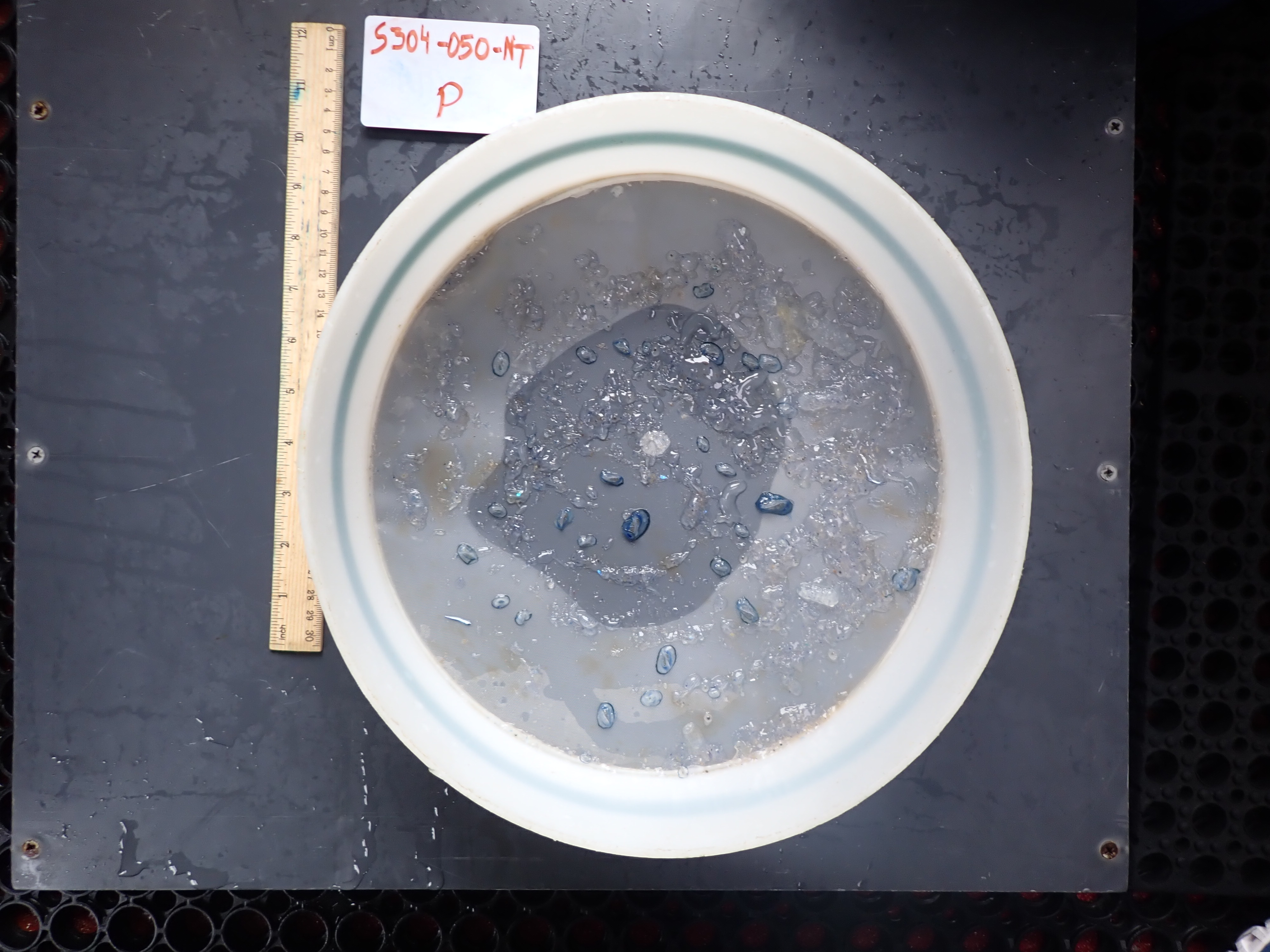

### HelmLab_S304-050-NT_07202022_Rinse.tif

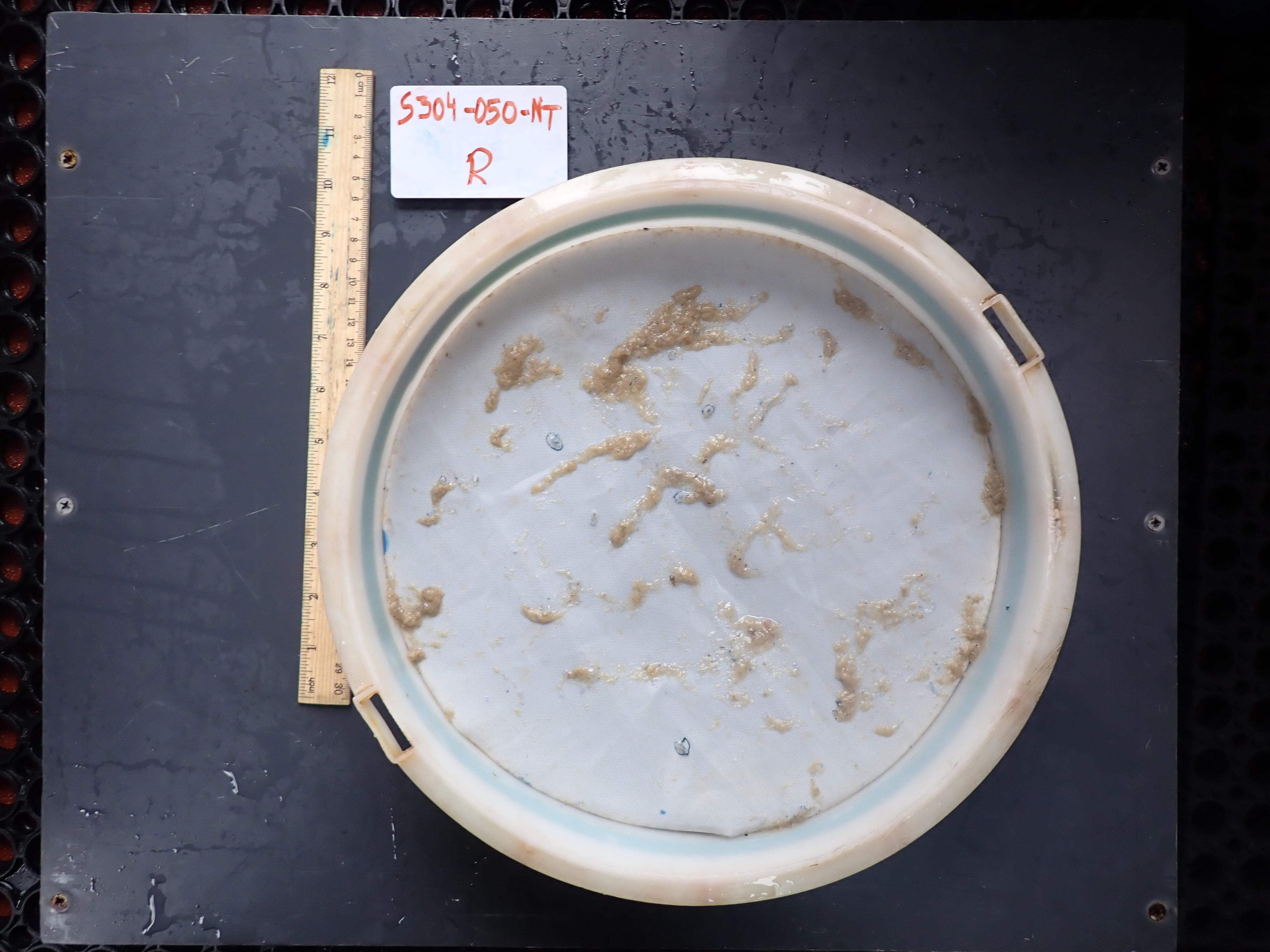

### SJR_019_screen.tif

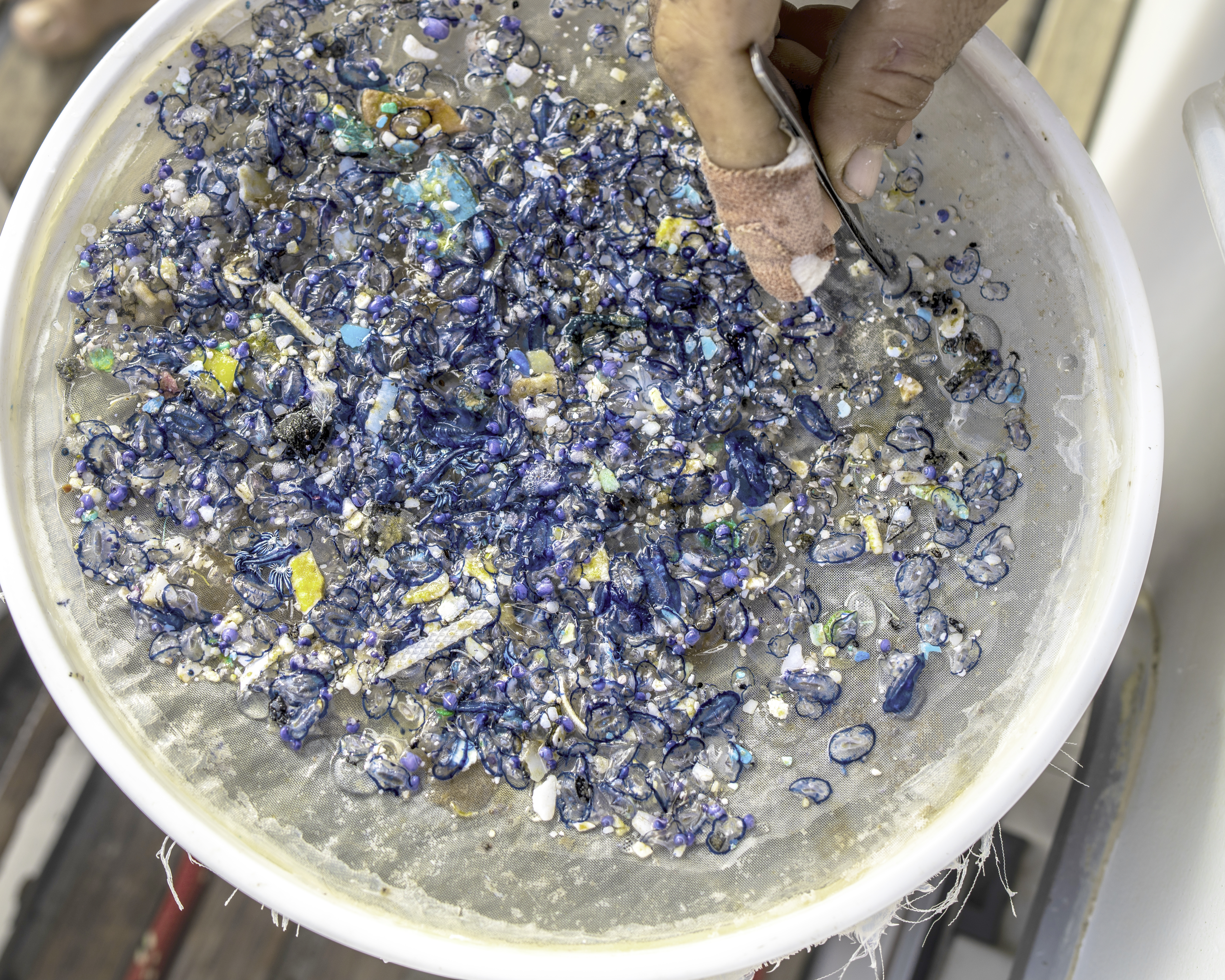

### SJR_31.tif

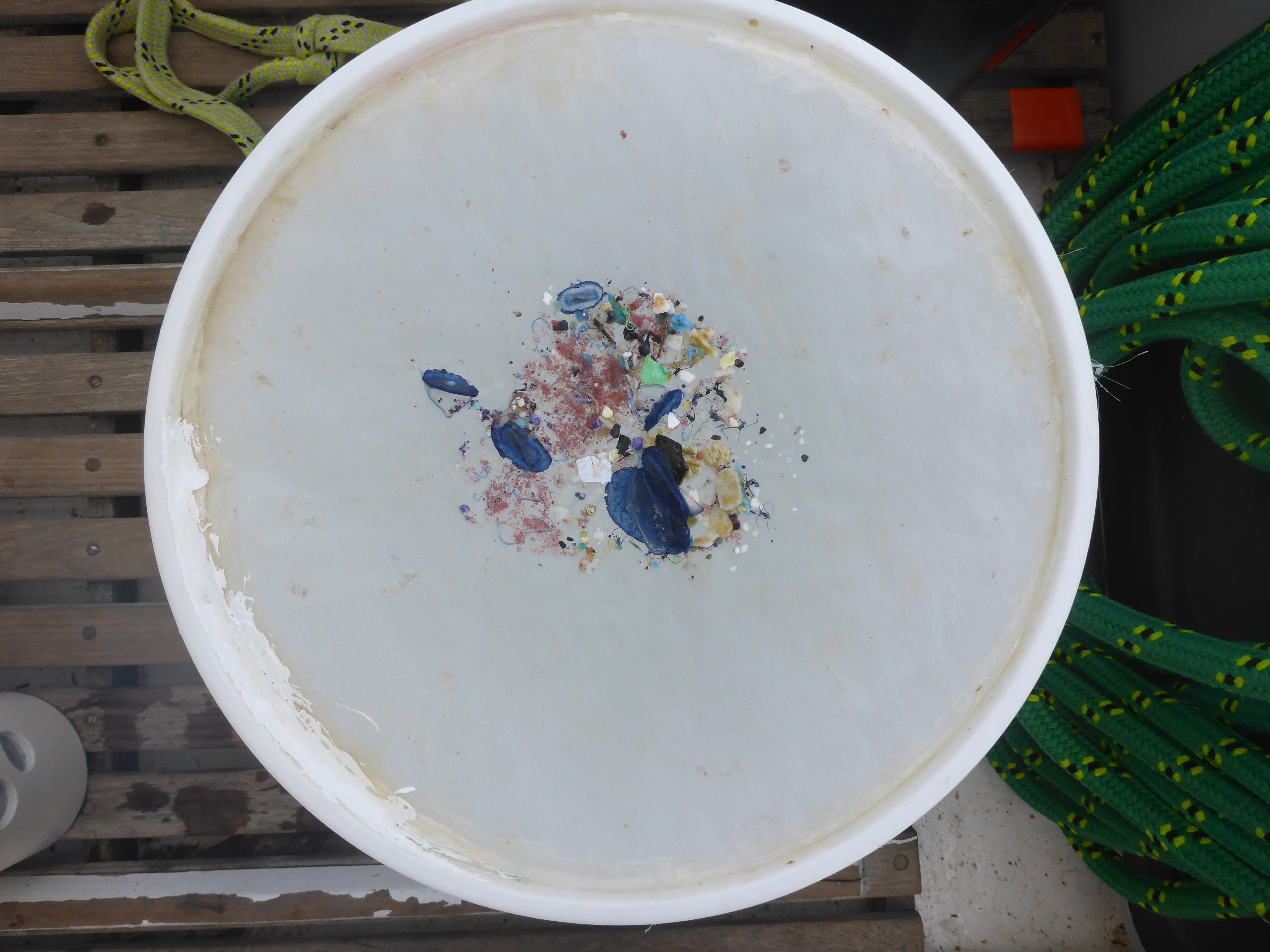

### SJR_34.tif

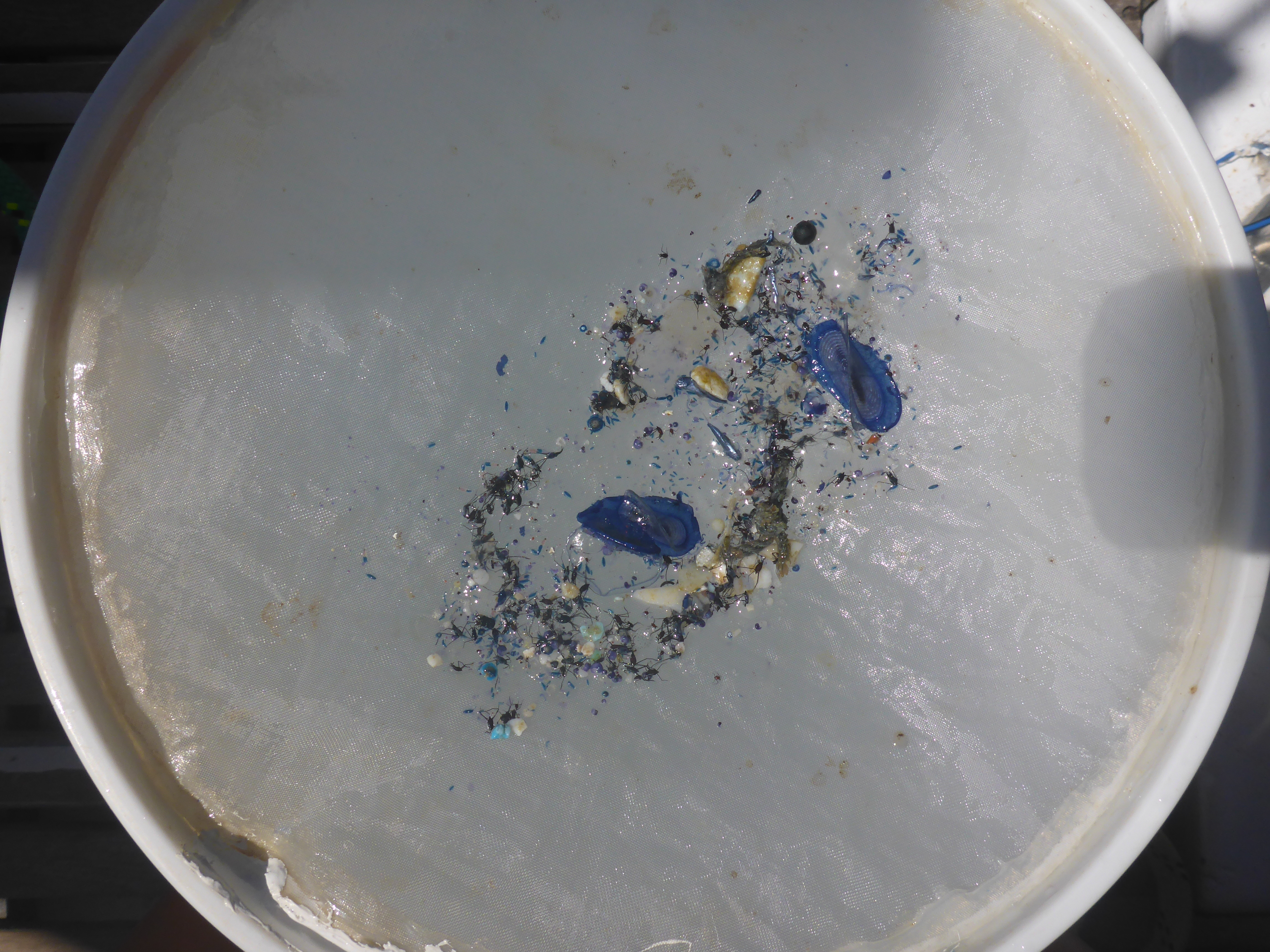

### SJR_37.tif

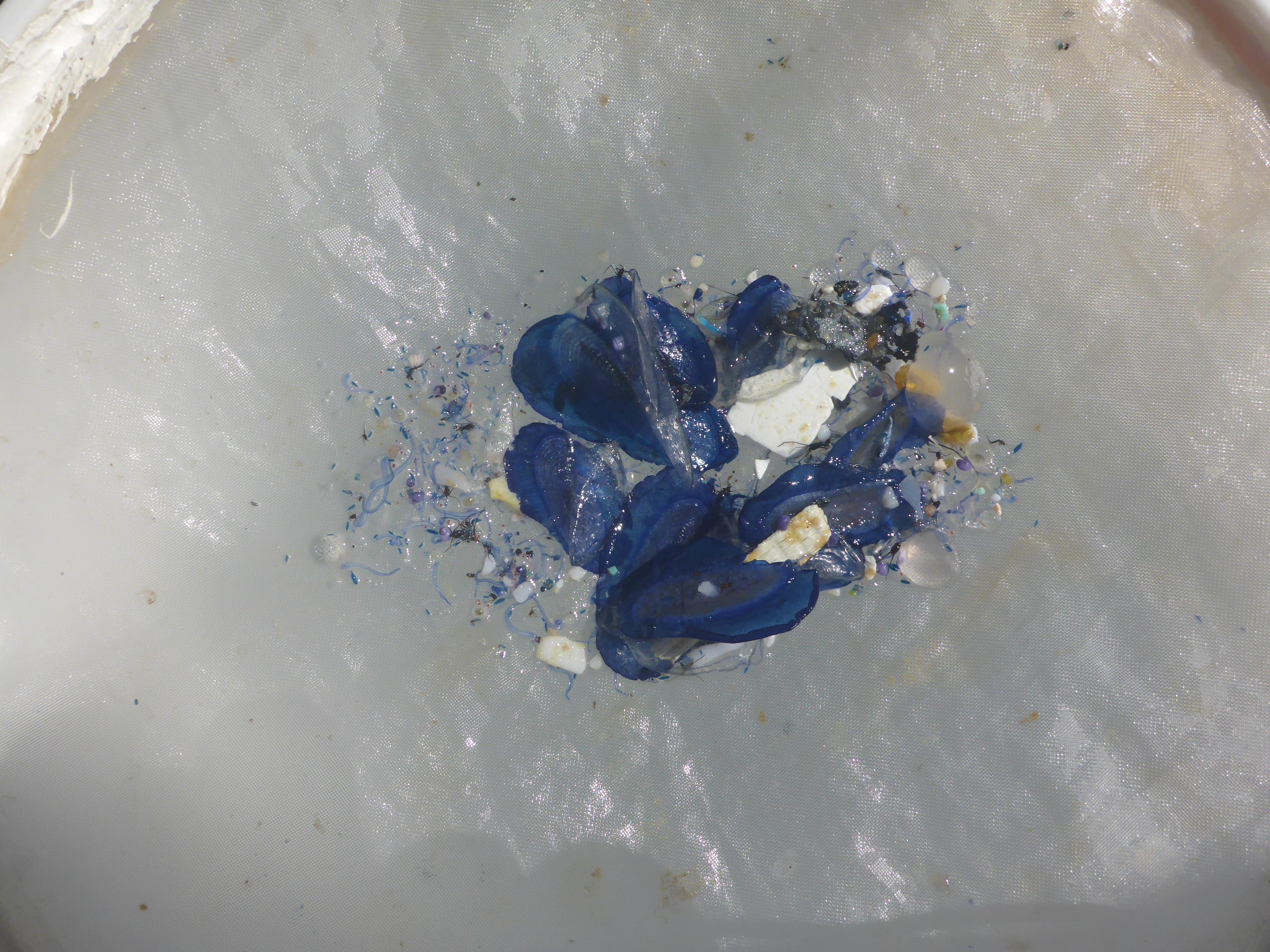

### SJR_38.tif

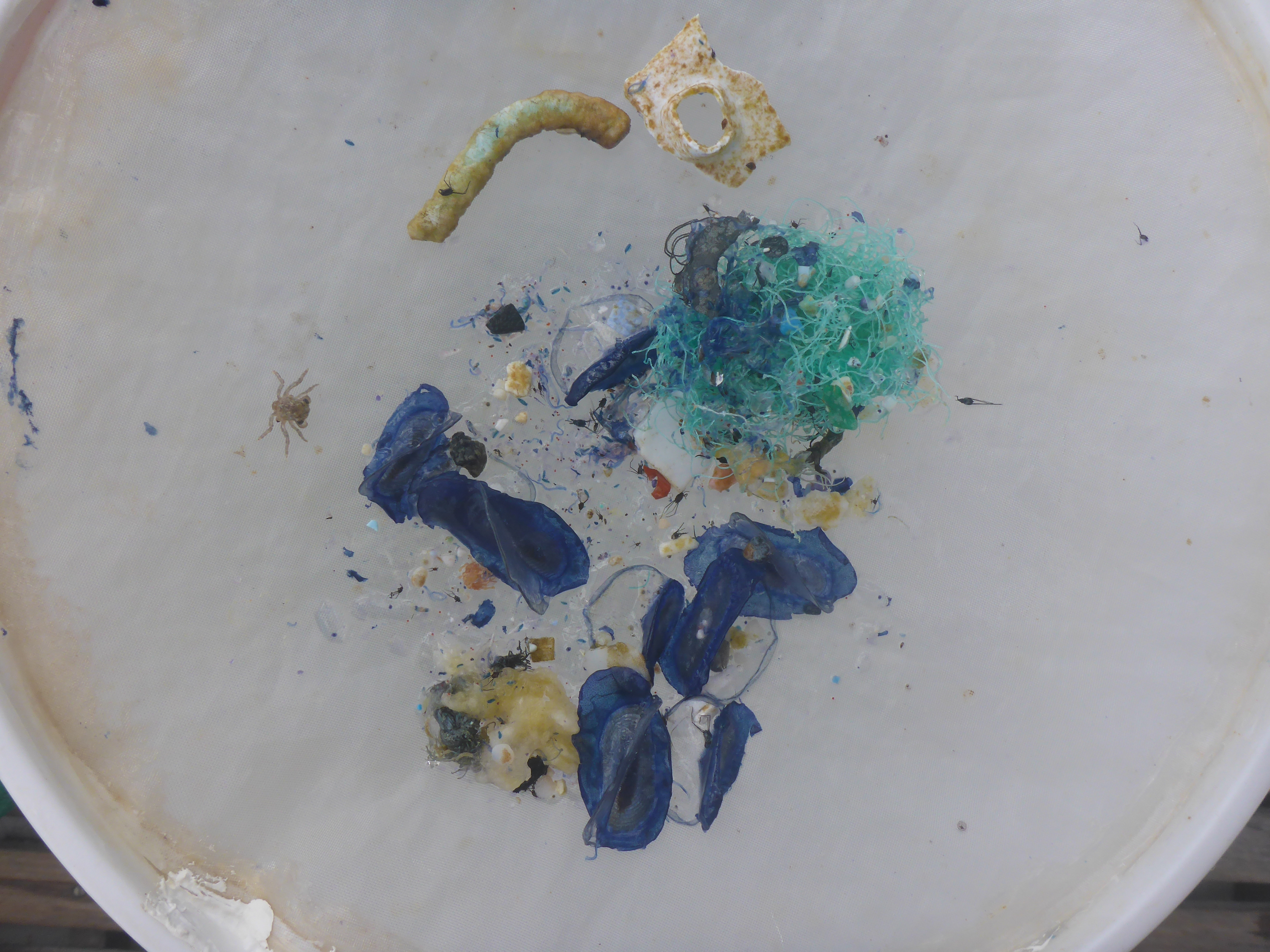

### Supp File C: mitochondrial ML tree

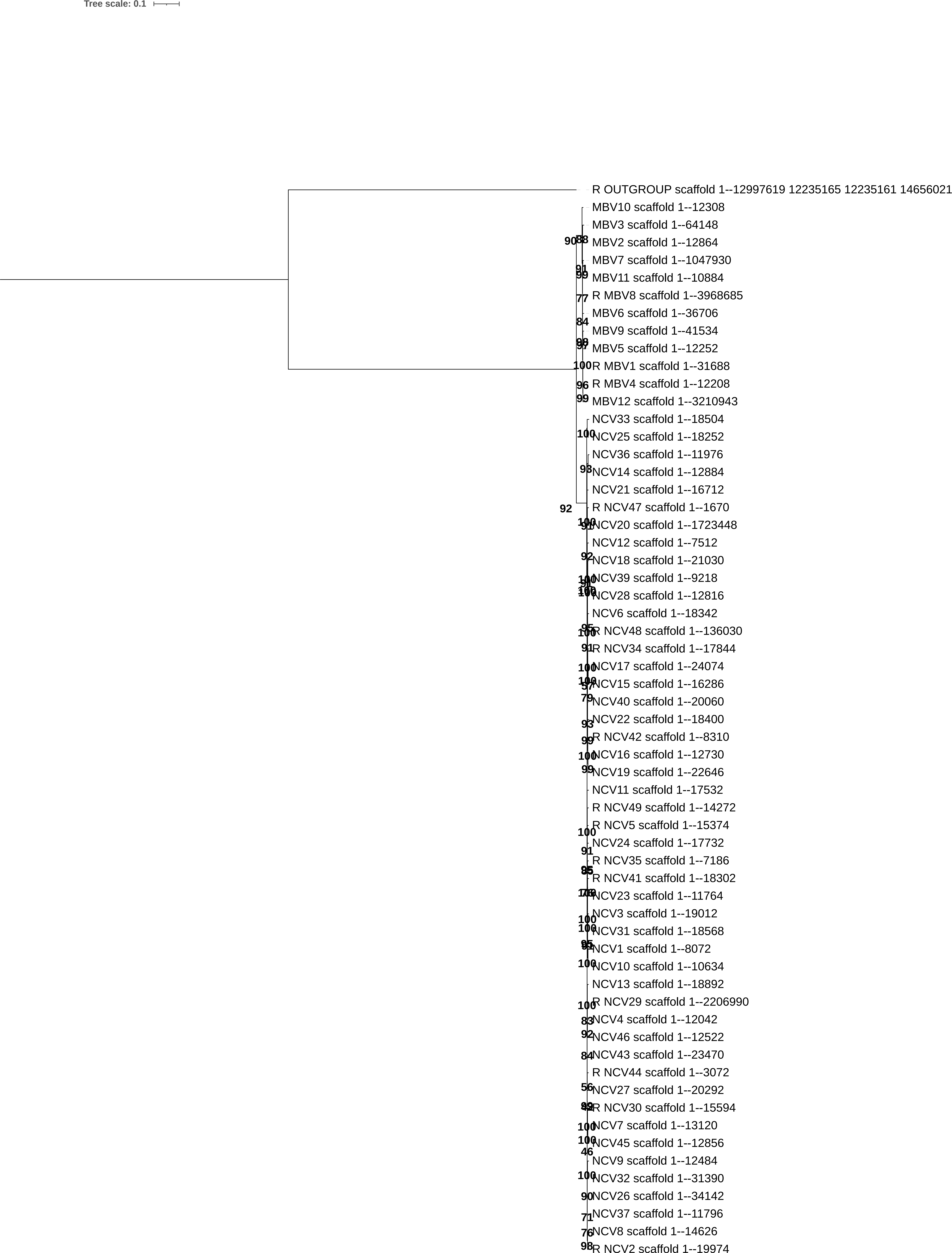

### TLS_112.tiff

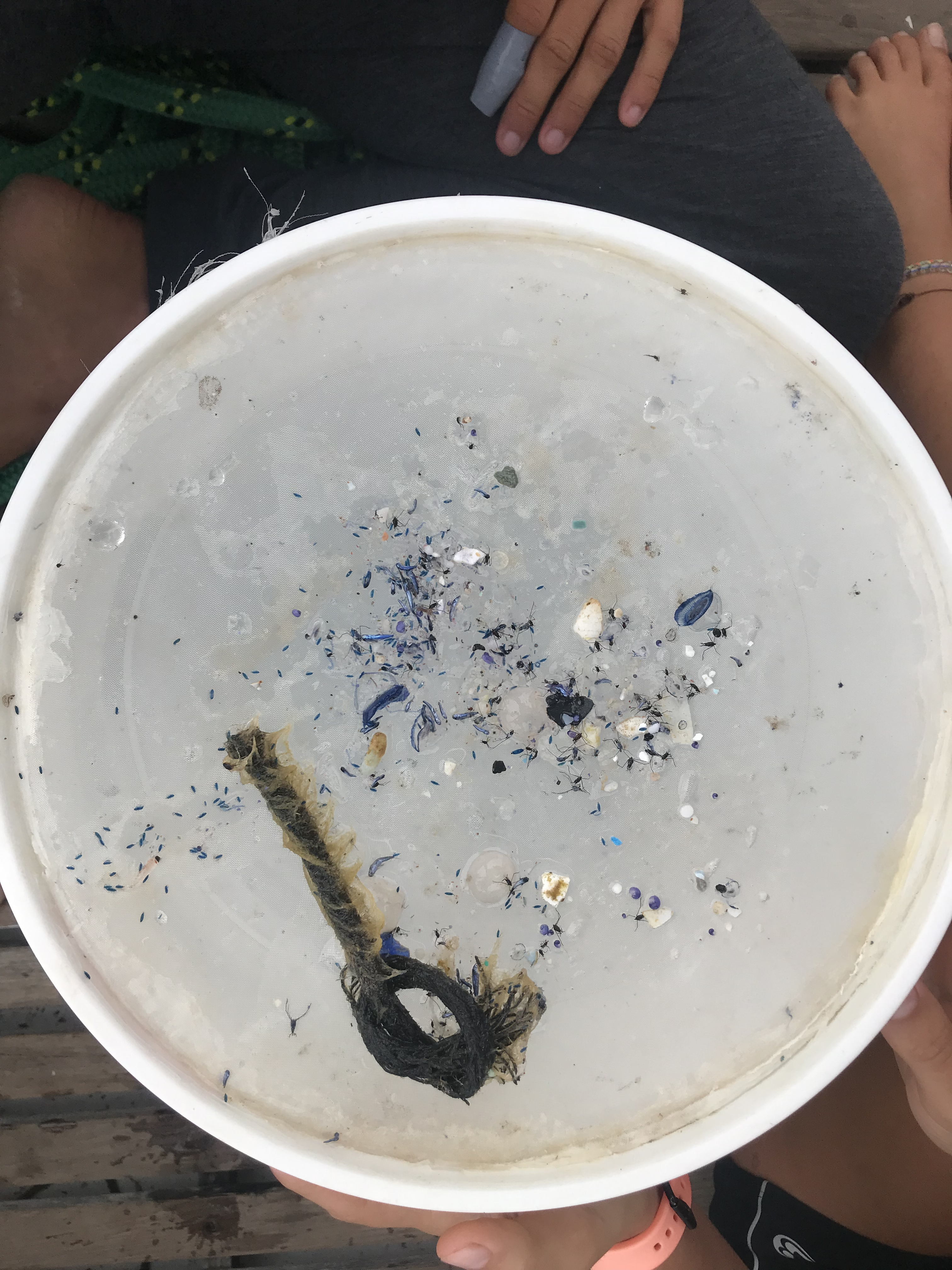

### TLS_134.tif

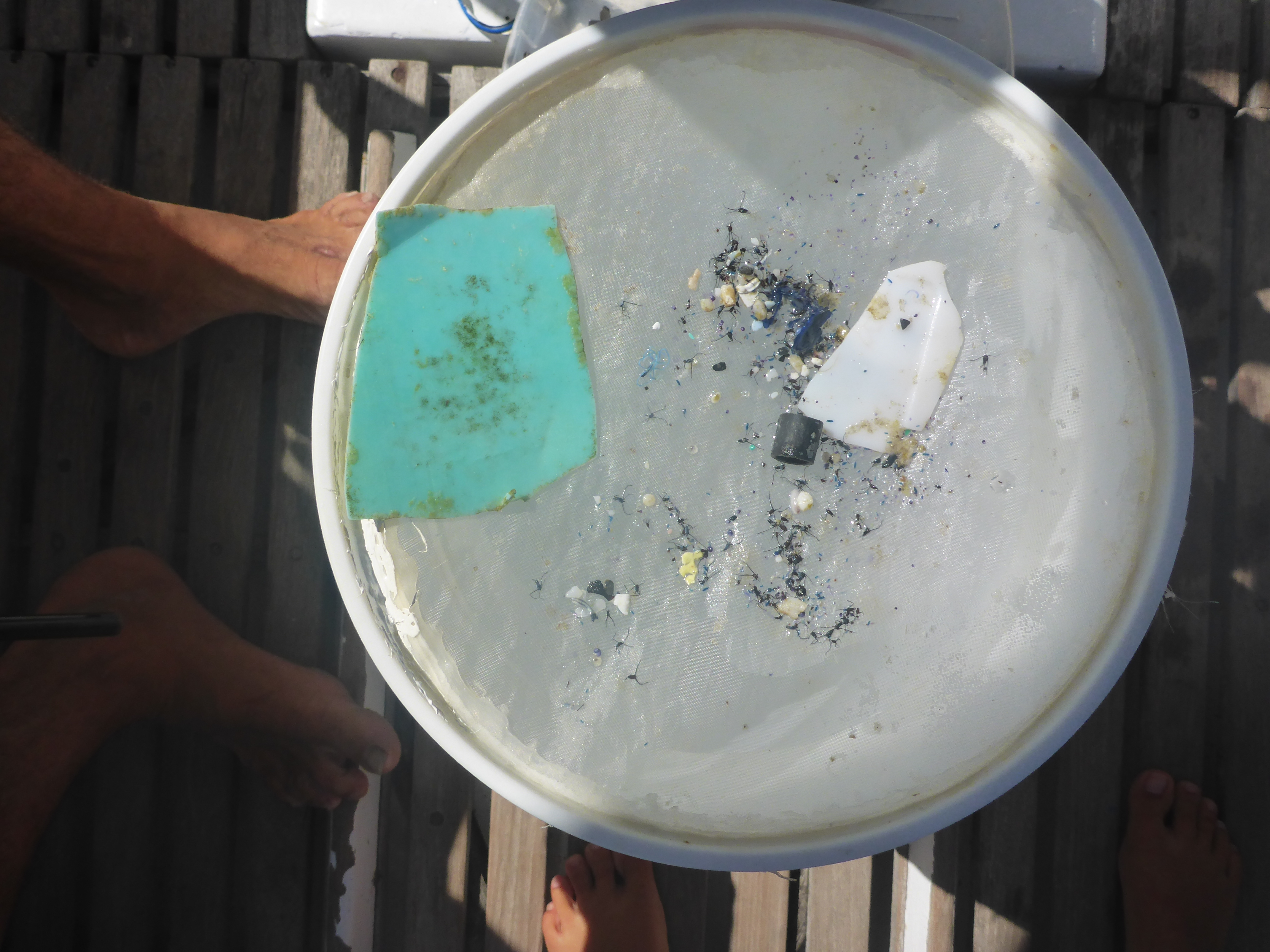

### TLS_135.tif

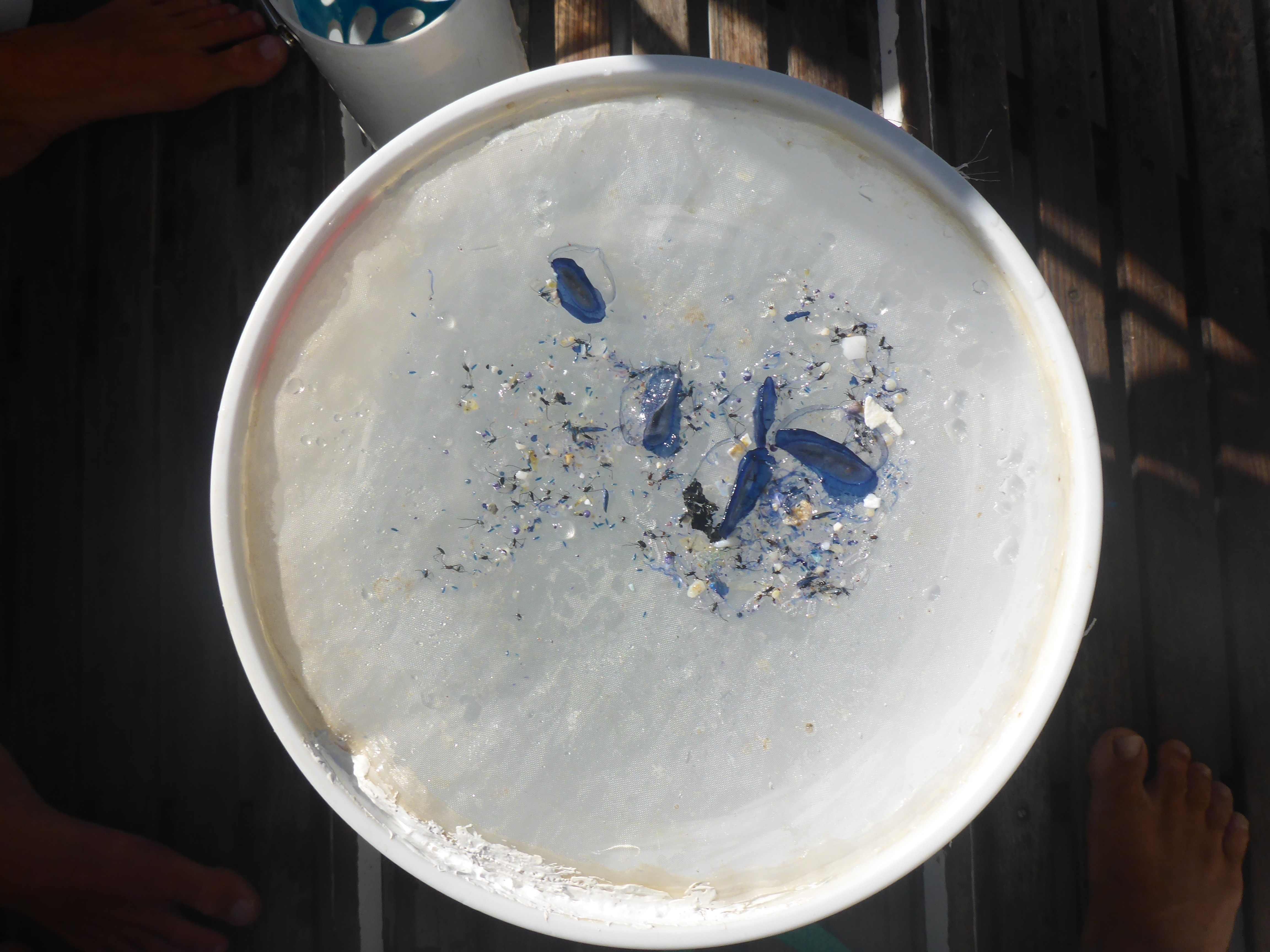

### TLS_138.tif

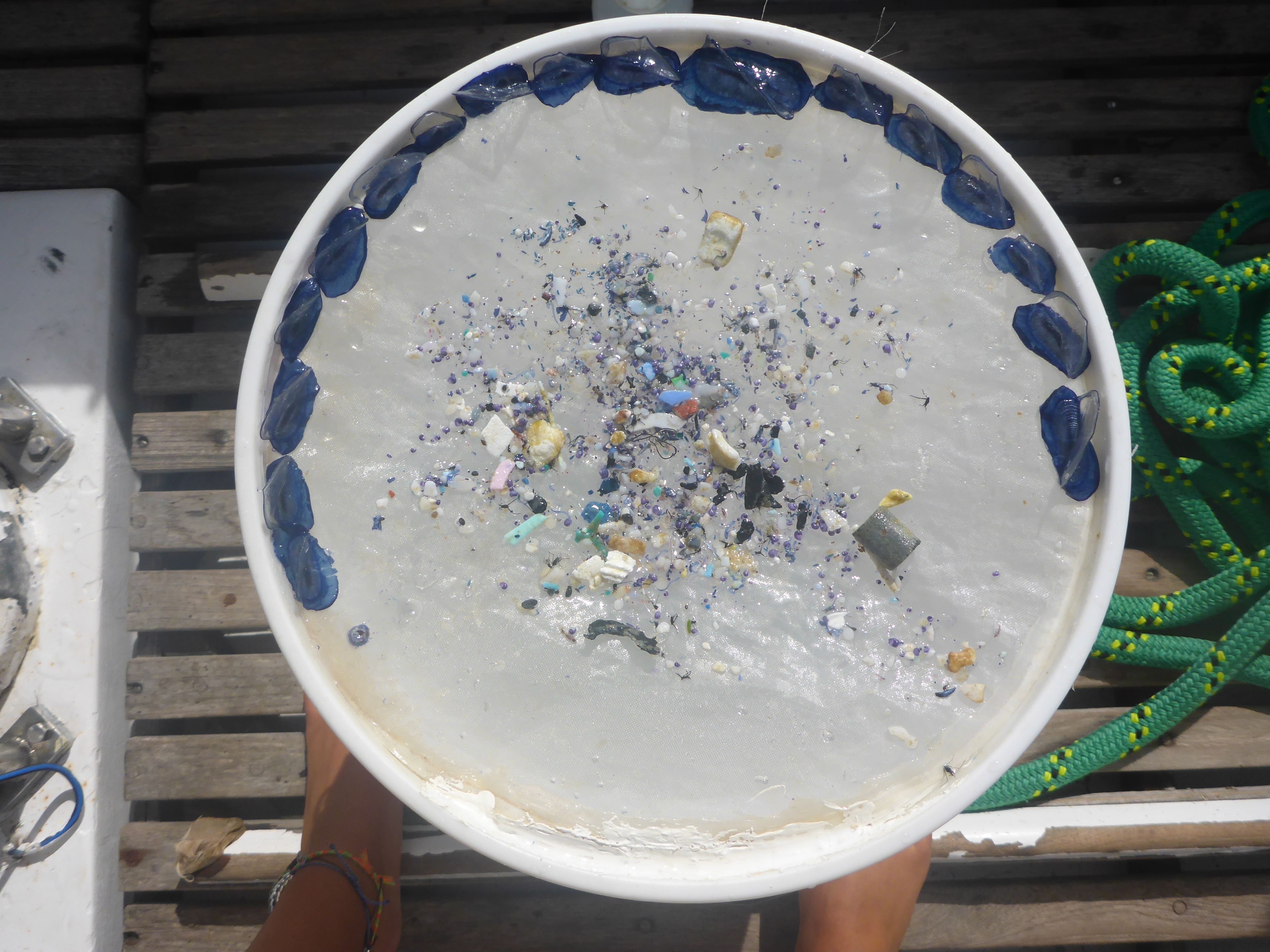

### VS_03.tif

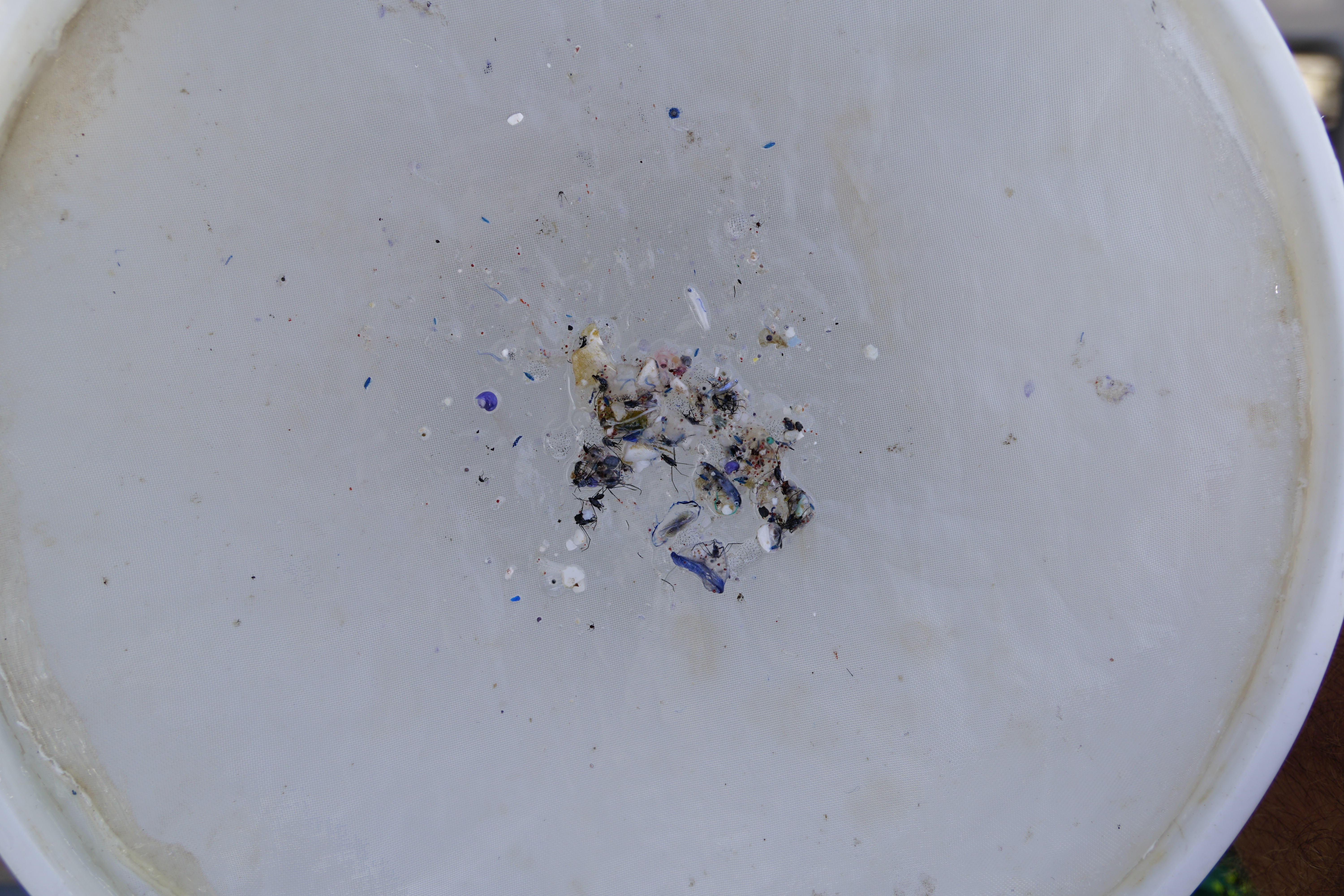

### VS_09.tif

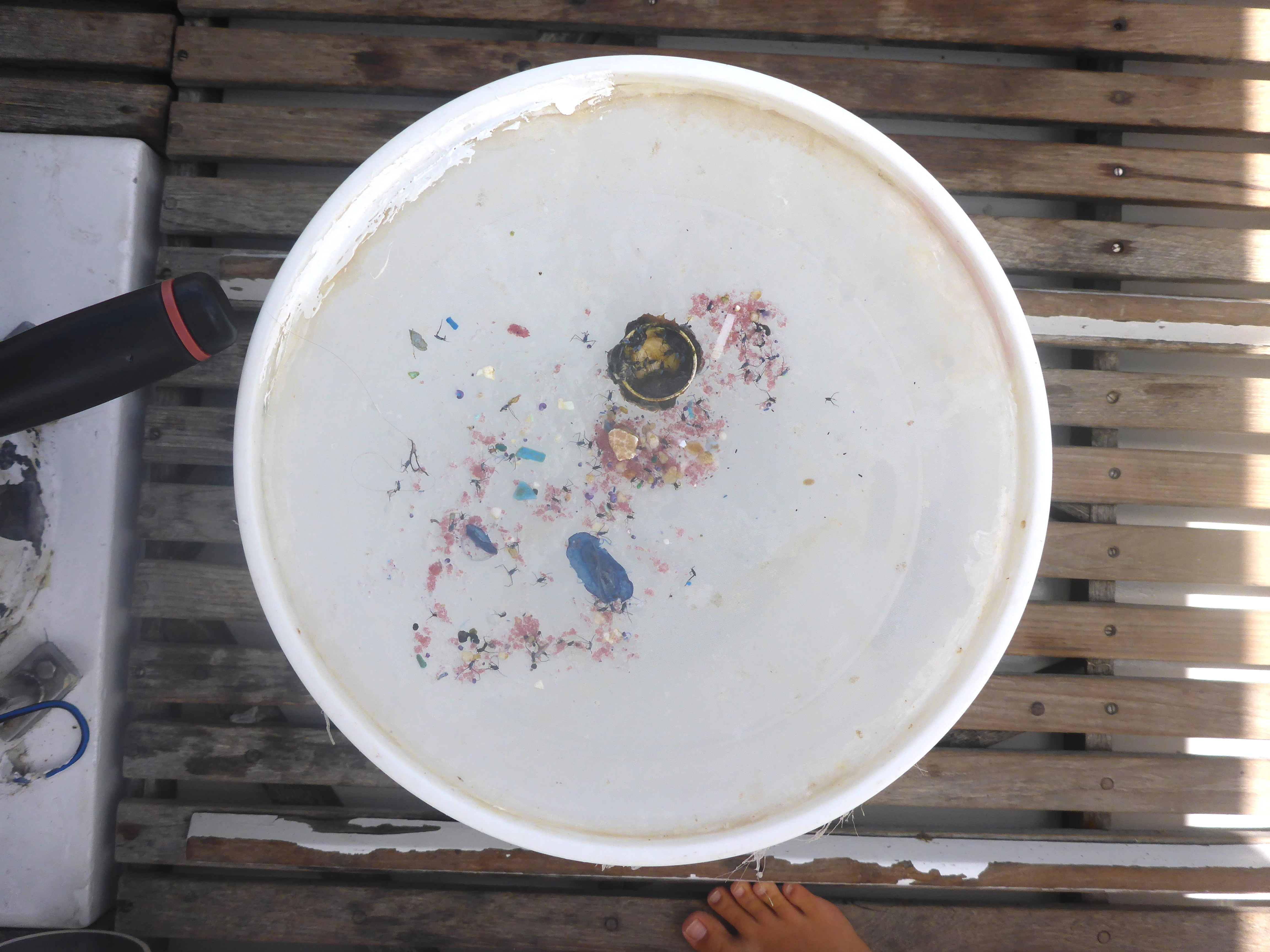
